# Protonated Structure of EmrE Reveals C-terminal Tail Gating Mechanism

**DOI:** 10.64898/2026.05.28.728512

**Authors:** Ashley B. Hiett-Borcik, Benjamin D. Harding, Merissa Brousseau, Owen Warmuth, Collin G. Borcik, Chao Wu, Eva Uhlemann, Claudia C. Cornilescu, Grace Reichert, Chad M. Rienstra, Katherine A. Henzler-Wildman

**Affiliations:** University of Wisconsin-Madison, Department of Biochemistry (Madison, WI); National Magnetic Resonance Facility at Madison, University of Wisconsin-Madison (Madison, WI); University of Wisconsin-Madison, Department of Chemistry (Madison, WI); Morgridge Institute for Discovery, University of Wisconsin-Madison (Madison, WI)

**Author notes:** Corresponding Author: Correspondence to Katherine A. Henzler-Wildman. Current affiliations - Eva-Maria Uhlemann, Vaccine and Infectious Disease Organization (VIDO), University of Saskatchewan - Chao Wu, Institute of Systems and Physical Biology, Shenzhen Bay Laboratory, Shenzhen 518132, China - Claudia C. Cornilescu, Division of Hospital Medicine, Stanford University School of Medicine, Stanford, California - Collin Borcik, Promega Corp. - Benjamin Harding, Resynant - Merissa Brousseau, University of Wisconsin-Madison, School of Medicine and Public Health, Department of Pathology and Laboratory Medicine.

## Abstract

The multidrug efflux pump EmrE is one of the smallest known active transporters and has become a model system for studying multidrug recognition and transport. While recent high-resolution structures have illuminated its dynamic substrate binding pocket, the conformations of its interhelical loops and C-terminal tail, regions critical for controlling proton coupling and gating, remain poorly characterized. Here, we report the high-resolution structure of protonated S64V EmrE determined using solution and solid-state NMR data. This new structural model shows the C-terminal tail occluding the open face of the transport pore, providing a structural basis for how EmrE minimizes proton leak in the absence of substrate. These findings support growing evidence that relatively simple model transporters must leverage an occluded state during alternating access to avoid physiologically unfavorable proton leak.

## INTRODUCTION

Bacteria confer multidrug resistance through multiple mechanisms including efflux, where toxic compounds are expelled from the cytoplasm through membrane-embedded transporter proteins.^1,2^ The majority of bacterial multidrug efflux pumps couple the expulsion of drug to the import of proton along the cellular proton-motive force (PMF).^3^ Such active efflux of drug against its electrochemical potential requires that the transporter must always remain closed on at least one side of the membrane.^4^ To achieve this, many ion-coupled transporters pass through a transient, fully-occluded intermediate state in the transition between open-in and open-out states.^5,6^ Structural characterization of several families of larger transporters has revealed multiple gating mechanisms where a part of the protein occludes substrate accessibility at the open side of the transporter.^5,7^ The gate serves as a barrier between the substrate binding site and the external environment, preventing rapid equilibrium between free and bound substrate. The structural basis by which substrate binding is occluded, the conformational changes that occur to alternate accessibility of the binding site across the membrane, and how binding of substrate and driving ion affect the energy landscape of the transporter to regulate gate opening are key to understanding transport regulation and how secondary transporters couple transport of driving ion and substrate.

The *Escherichia coli* small multidrug resistance (SMR) transporter EmrE is one of the smallest known multidrug efflux pumps and has been widely used as a model system for studying proton coupled multidrug efflux.^8,9^ Structurally, EmrE consists of two sequentially identical subunits, each a 110-amino acid monomer with four transmembrane (TM) helices, oriented topologically as an asymmetric homodimer.^10^ Two glutamate residues (Glu-14_A,B_) are embedded within the central binding pocket and are essential for binding and transport of drug and proton across the membrane. To confer multidrug resistance, the two subunits of EmrE exchange conformations to alternate access of the central binding pocket and antiport substrate and proton across the bacterial membrane.

EmrE utilizes different transport modes depending on substrate identity, with experimentally confirmed 2:1 and 1:1 H⁺/drug⁺ antiport and drug-gated proton leak.^11^ This mechanistic flexibility is incorporated within the free-exchange model of EmrE transport^12^, a universal-exchange model including the minimum states of EmrE observed with NMR and other biophysical methods under near-physiological pH and temperature. The free-exchange model^12^ predicts that uncoupled proton leak through EmrE should be the most kinetically favorable mode of transport in the absence of substrate. However, EmrE exhibits minimal proton leak under a pH gradient in the absence of substrate.^13^ A recent study reconciled this discrepancy and demonstrated the C-terminal tail of EmrE plays an essential role in gating proton leak.^14^ Deletion of the four C-terminal residues (Δ107-EmrE) results in significant uncoupled proton leak through EmrE, demonstrating that the intact tail is required to maintain tight regulation of proton flux.^15^ Moreover, the protonation state of H110 is allosterically linked to drug binding at Glu-14 in the core of the transporter, further implicating the tail in the coupled transport cycle of EmrE.^16^

Prior studies also implicated the C-terminal tail and adjacent loop regions of EmrE in transport function. Scanning mutagenesis assays demonstrated that mutations to the TM 3-4 loop residue Asp-84 impairs drug resistance in bacteria,^17,18^ and mutating Asp-84 alongside TM 1-2 loop residue Glu-25 severely impairs resistance in bacteria and diminishes EmrE transport activity *in vitro*.^18^ An additional study found that the substrate TPP^+^ transiently binds at a secondary peripheral site in addition to binding in the core of the transporter at Glu-14, and postulated substrate-loop interactions at Glu-25 and Asp-84.^19^ Furthermore, recent work has shown that certain substrates trigger uncoupled proton leak by perturbing C-tail and loop residues rather than engaging the canonical drug-binding pocket.^20,21^

Despite the demonstrated functional importance of the C-terminal tail and loop regions of EmrE, these regions remain poorly defined in existing structural models. Current experimentally derived models of EmrE from X-ray crystallography^22^ and NMR spectroscopy^23–25^ provide valuable insights into drug binding in different protonation states, the similarities and differences in the interactions of EmrE with diverse substrates in the primary binding site in the transport pore, and conformational changes upon ligand engagement. These structural models all reflect the same long-established arrangement of the transmembrane helices^26^, but fail to resolve the conformation of the interhelical loops and the C-terminal tail—regions now known to modulate proton leak and transport activity. For example, the highest resolution crystal structures of both substrate-free and substrate-bound EmrE^22^ (PDBs 7MH6, 7MGX, 7SVX, 7SSU, 7SV9, 7T00) were solved in complex with two L7 monobodies that bind directly to the first interhelical loop, and the protein was engineered with three mutations in the same loop (E25M, W31I, V34M) to facilitate crystallization. These crystallization conditions and mutations influence the native interhelical loop conformation and therefore limit functional interpretation in these regions (Fig. 1A). The ^19^F solid-state NMR structures of TPP^+^ bound EmrE at low pH^27^ (PDB 7JK8) and high pH^23^ (PDB 7SFQ) were determined using distance restraints between a fluorinated substrate and protein amides, a technique which is effective for distances up to about 20 Å. As the interhelical loops and C-tail are located farther from the primary substrate binding site at Glu-14 and are largely outside of the measurable distance range, those regions were poorly restrained or unrestrained (Fig. 1B). Similarly, the NMR structure of protonated EmrE^25^ (PDB 8UWU) includes only three NOE restraints and 227 PRE restraints in the interhelical loops and C-terminal tail, and one EmrE subunit includes loop mutation L51I to diminish conformational exchange (Fig. 1C). The NMR structure of TPP^+^ bound heterodimer E14Q-EmrE/WT EmrE^25^ (PDB 8UOZ) is the most complete structure of EmrE with a larger number of structural restraints within the interhelical loops (Fig. S1), but has minimal restraints in the C-terminal tail. This leaves the structural basis for proton-gating by EmrE uncharacterized. Well-restrained structural data for the C-terminal tail and interhelical loop conformations of EmrE in substrate-free conditions is essential to understand both proton gating in the absence of substrate and the transition between coupled and uncoupled transport by this very small transporter.

**Figure 1.**
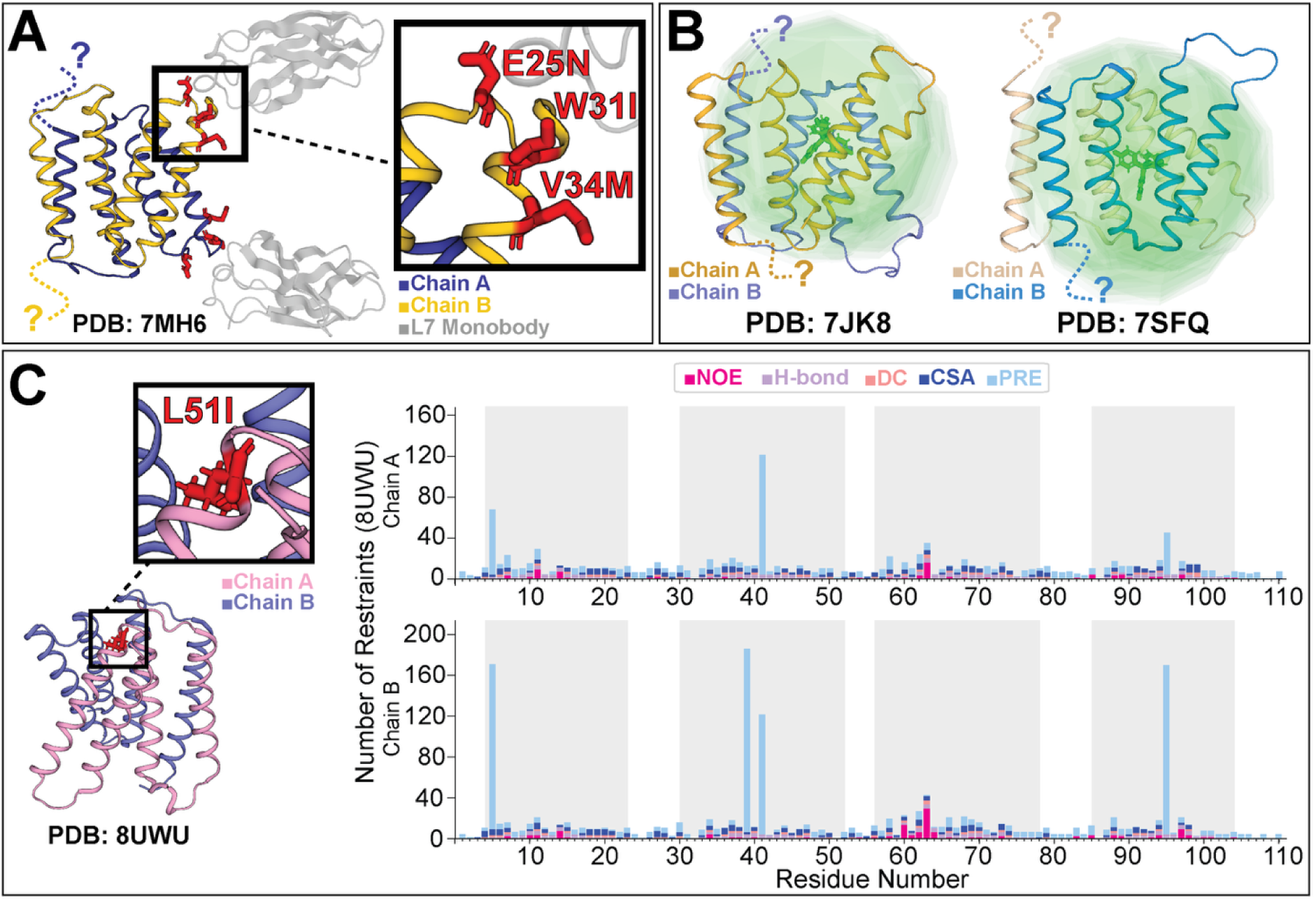
Current experimental structures of EmrE have limited resolution or mutations in loop and C-terminal tail regions. A) The highest resolution crystal structures of EmrE include three loop mutations and do not model six C-terminal tail residues. B) NMR structures of F-TPP+ bound EmrE include distance restraints within a 20 Å radius from the substrate (shown as green orbs) which do not include peripheral loop residues. Five C-terminal tail residues are also excluded from these models. C) NMR structure of protonated EmrE (PDB 8UWU) includes loop mutation L51I and has fewer restraints in the loop and C-tail regions of EmrE (helices shaded in grey).

In this study, we use solution- and solid-state NMR data to determine a high-resolution structural model of lipid-embedded, low pH and drug-free S64V EmrE with a focus on revealing the structural basis of EmrE C-terminal tail gating. We selected these conditions because at this pH, NMR data shows both Glu-14 residues of the transporter are protonated, corresponding to a key step in the EmrE transport cycle.^12^ Furthermore, the S64V mutation in TM helix 3 minimally disrupts the structure and substrate binding affinity but slows the rate of conformational exchange, allowing site-specific NMR assignments without perturbing the loop or C-terminal tail regions.^17^ Our model is the first experimental structure to show the C-terminal tail interacts electrostatically with loop residues to occlude the transport pore and block access to the canonical binding pocket centered at residue Glu-14.

## RESULTS

### Structural modelling and Refinement statistics

Here, we present a structural model of substrate-free, protonated S64V-EmrE (Fig. 2A) determined using 6,202 restraints from both solution and solid-state NMR (Fig. 2B). 1,160 of the restraints (18.7%) are in the interhelical loops or C-terminal tail (Fig. 2C-D), providing significantly more data to restrain this region of the protein than prior structures. Importantly, our EmrE construct also has the WT sequence in these regions. Initial models were generated with Xplor-NIH^28,29^ using distance restraints, torsion angles determined by TALOS-N,^30^ and SSNMR water accessibility potentials^31^, with the overall fold initially restrained by the previously published crystallographic density of protonated EmrE.^22^ This enabled correct folding of EmrE into its well-known helical topology along the membrane normal, a fold that is observed across all prior structures in all lipid and detergent environments across all SMR variants for which structures have been determined (EmrE and Gdx).^22,23,25,27,32–35^ The ten structures with the lowest energy terms were integrated into explicit POPC lipid bilayers using CHARMM-GUI^36^ and further energy-minimized with NMR restraints using GROMACS^37^ (no restraints to crystallographic data were used at this stage). Refinement statistics are shown in Table 1. An average of 94% ± 0.6% of 5,692 total distance restraints (evaluated with 0 Å tolerance from input restraints) and an average of 91% ± 0.9% of 510 total torsion angle (ϕ, ψ, χ) restraints (evaluated with 0° tolerance from input restraints) are satisfied across the model bundle. Distance restraints cover the full extent of the sequence of both chains of EmrE, and no residue shows a disproportionally high number of violations (Fig. S2). PROCHECK^38^ analysis shows 94.7% ± 0.8% of residues in the most-favored Ramachandran space, 4.5% ± 0.8% in allowed regions, 0.6% ± 0.6% in generous regions, and 0.28% ± 0.4% in disallowed regions. MolProbity^39^ reflects a clash score of 0 for all models and an average overall score of 0.961 ± 0.22. Our structural model is the first to show the C-terminal tail of EmrE adopts a consistent structure covering the open face of the drug-binding pocket, occluding access to the binding pocket in the absence of substrate.

**Figure 2.**
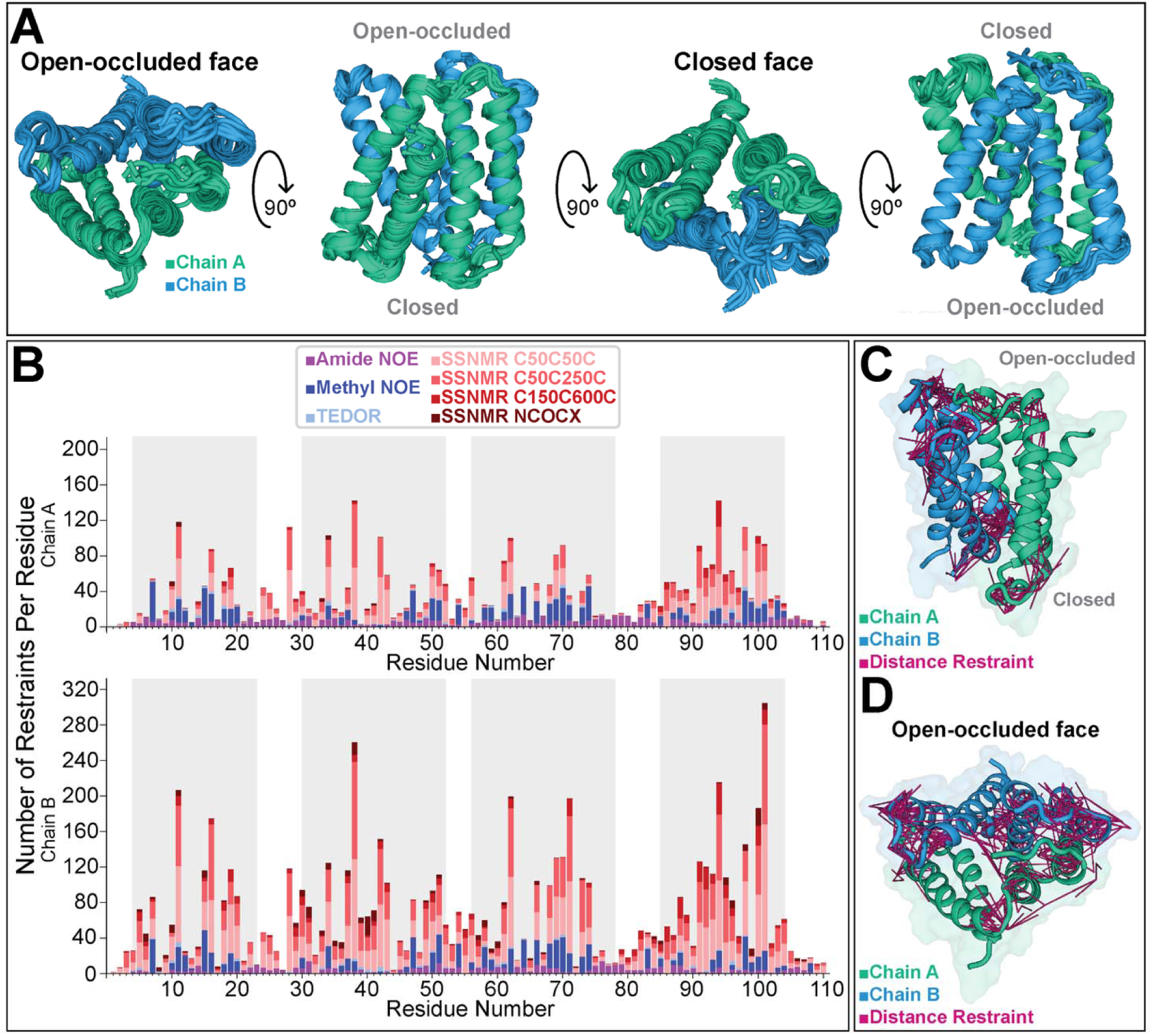
Low pH EmrE structure. A) 360° view of the 10 lowest energy structures of low pH S64V EmrE. B) Bar graph quantifying reciprocal NMR-derived restraints per residue of EmrE. Distance restraints span the entire transporter and include over a thousand distance restraints within the loops and C-terminal tails (helices shaded in grey). C) Side view and D) Top view of EmrE showing the distance restraints that constrain the loops and C-terminal tail.

**Table 1.**
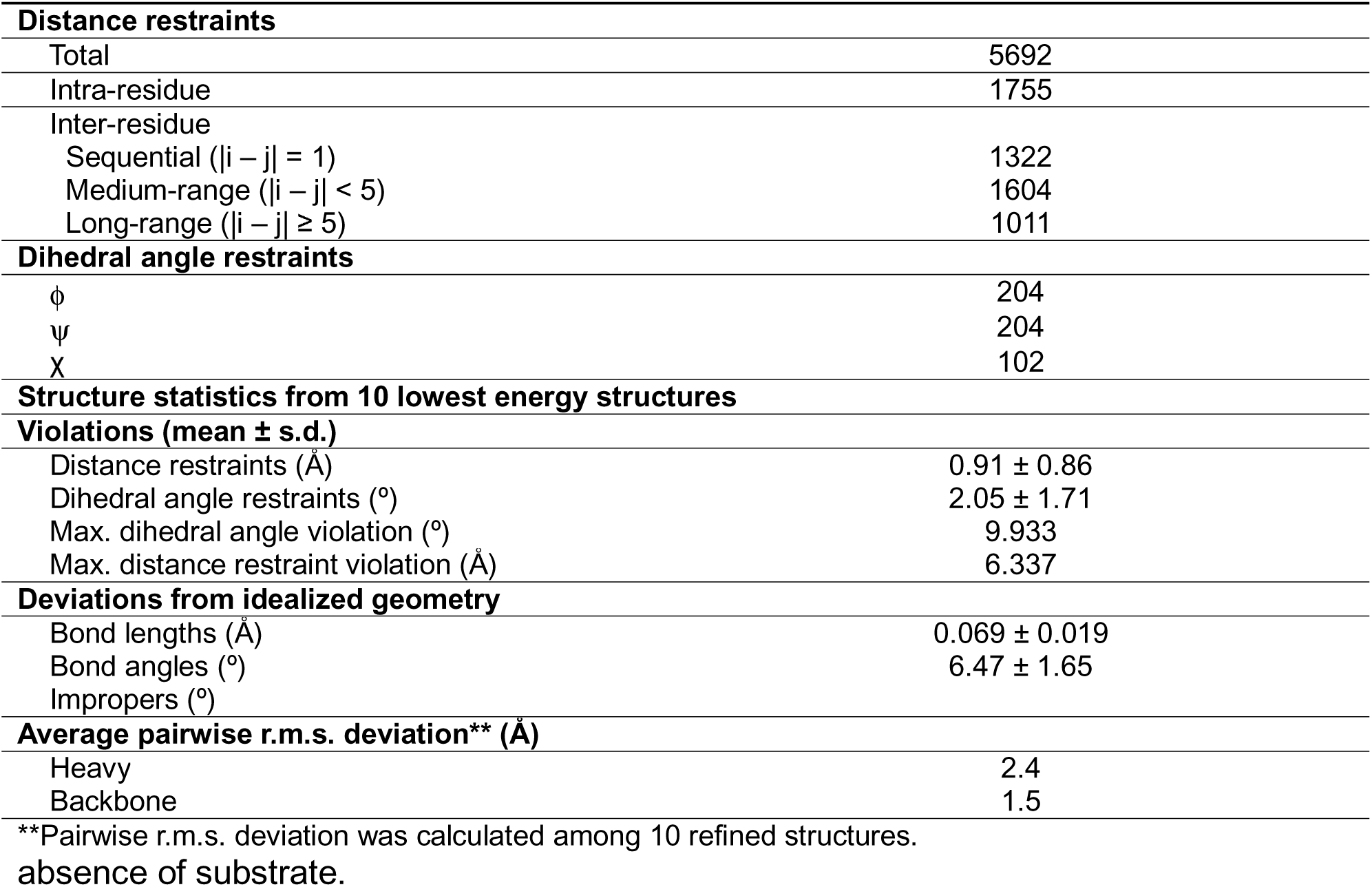
NMR distance and dihedral restraints and refinement statistics for S64V-EmrE pH 5.8.

### Electrostatic interactions coordinate C-terminal tail gating

Our structure (Fig. 2A) is consistent with recent work showing the C-terminal tail of EmrE acts as a secondary gate against uncoupled proton leak through the substrate-free transporter.^14^ Previous studies indicate that the C-terminal tail becomes displaced upon drug binding to the primary binding site at Glu-14.^14,16,25^ This suggests disruption of EmrE loop-tail interactions should not significantly impair drug/proton antiport and the ability of EmrE to confer resistance to toxic compounds. Extensive scanning mutagenesis studies of EmrE found only a few minor impacts on EmrE-mediated bacterial resistance or *in vitro* drug transport upon mutation of TM loop or C-terminal tail mutations,^17,18,40^ and C-tail truncation (Δ107-EmrE) did not impair ethidium resistance relative to WT-EmrE.^14^ This led to underappreciation of the functional importance of the C-terminal tail. However, mutation of loop or C-terminal tail residues or truncation of the C-terminal tail in the absence of drug substrate leads to enhanced proton leak sufficient to cause diminished growth in bacteria.^14,21,41^

Recent molecular modeling of EmrE in lipids with and without the C-terminal tail suggested that coordination between electrostatic residues in the loops and C-terminal tail of the open side of EmrE led to occlusion of the fully protonated transporter, which would prevent proton release from Glu-14 and transport in the absence of drug-substrate.^14^ Within this model, key interactions included a salt bridge between residue Asp-84_B_ and Arg-106_A_ and hydrogen bond between Thr-56_A_ and His-110_A_. These interactions are consistent with our experimental structural restraints, with Asp-84_B_ and Arg-106_A_ forming a salt bridge and Thr-56_A_ and His-110_A_ hydrogen bonding in several of our 10 lowest energy structures. Backbone order parameters (S^2^) estimated from the backbone chemical shifts using TALOS-N^30^, indicate that the loops and C-terminal tail are dynamic but not fully disordered (Fig. S3). This is consistent with the limited sensitivity of the tail in solid-state NMR cross-polarization experiments. However, the unique chemical shifts and peak intensities similar to other residues observed for the C-terminal tail in the solution NMR spectra indicates that this region is in an extended conformation and unique environment. The 10 lowest energy structures reflect this with a consistently extended structure for the C-terminal tail of chain A that occludes access on the “open” face of the transporter as described below, but vary in the precise conformation and interactions.

EmrE is an asymmetric antiparallel homodimer, with distinct structures for each subunit. In our structure, the “open” face of EmrE is gated by the C-terminal tail, which extends between the interhelical loops of the two subunits, occluding access to the central binding pocket at Glu-14 (Fig. 2A). In prior structures lacking restraints or resolution of the C-terminal tail, the “open” face of the transporter is fully open to water with clear access from the aqueous compartment to Glu-14. On the closed face of the transporter, the ends of the transmembrane helices and the interhelical loops come directly together to occlude access and the C-terminal tail is far more disordered than on the “open” face (Fig. 2A). In EmrE, alternating access occurs through a conformational switch where the two subunits of the homodimer (chain A and chain B) swap conformations^42^. This unique mechanism is only possible because of the antiparallel topology of the transporter. Calculation of the chemical shift differences between the two chains highlights the regions where the local structural and chemical environment is most distinct between the two subunits (Fig. 3A). We observe large chemical shift perturbations in the loop and C-terminal tail residues involved in the electrostatic interactions that coordinate the C-terminal gate and their close neighbors, including Lys-22, Thr-50, Thr-56, Leu-83, Arg-106 and Thr-108. Superimposing the chain A and B structures shows high structural similarity between the two chains for several of these residues (Fig. 3B), consistent with differences in the chemical environment, such as local electrostatics, as the most likely origin of the observed chemical shift differences. Plotting the chemical shift differences on the structure shows a network of residues with significant perturbations on the solvent-exposed faces of EmrE, including the C-terminal tail (Fig. 3C-D). This is consistent with different structural environment of the C-terminal tail residues in the two subunits of the homodimer, implicating their functionally distinct roles on either face of the transporter.

**Figure 3.**
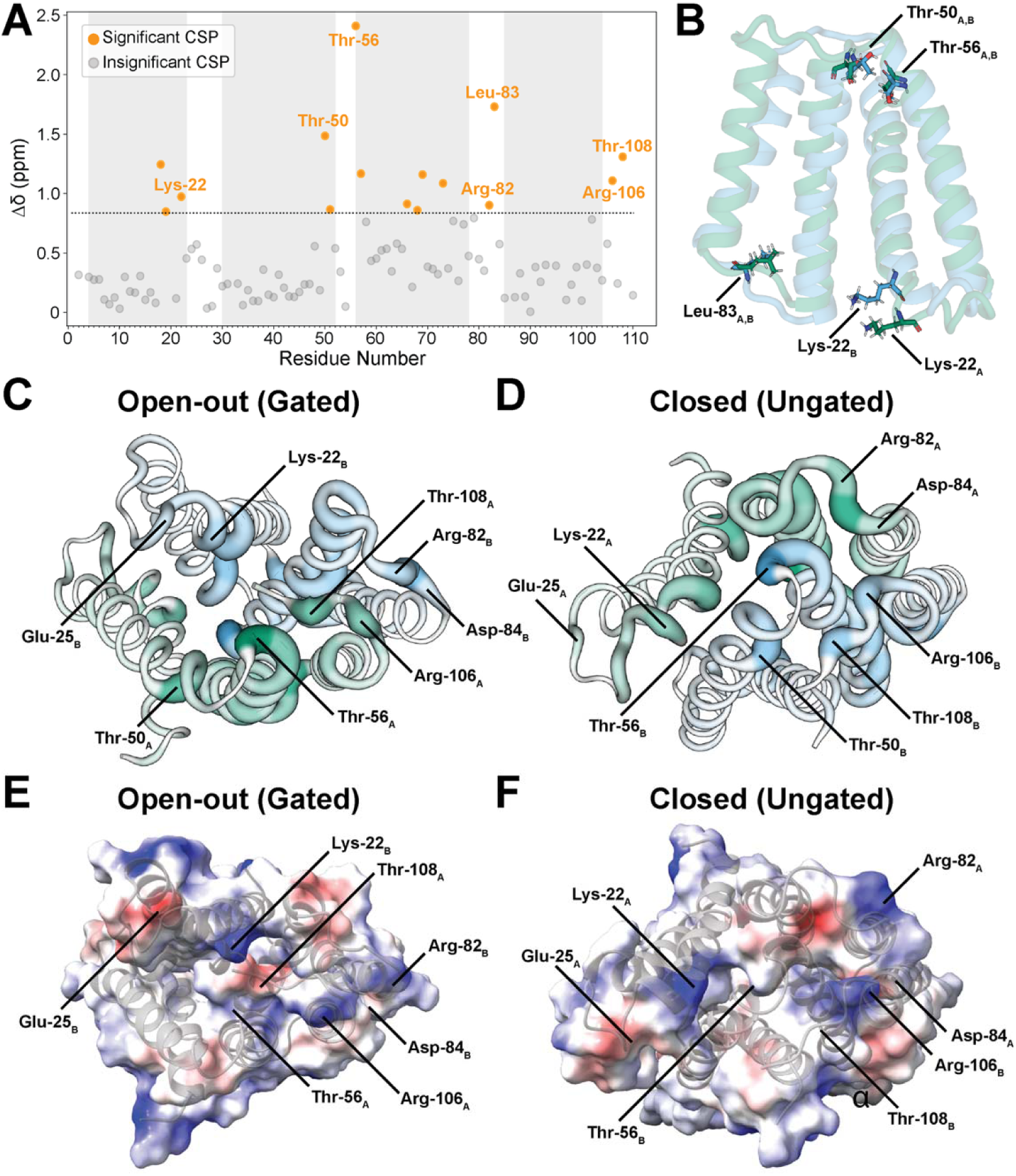
Chemical shift differences between the subunits highlight gating interactions. (A) The chemical shift difference (Δδ) between chains A and B of EmrE is plotted for each residue. The dashed line indicates the significance threshold of one standard deviation from the mean. (B) Superposition of chain A (green) and chain B (blue) shows the regions of structural similarity and differences between the two chains in the asymmetric homodimer. Large chemical shift differences are observed in some regions with high structural similarity. (C) The open-out face and the (D) closed face of EmrE with CSPs plotted as b-factors. (E) The open-out face of EmrE shows electrostatic interactions which may coordinate the C-terminal gate. (F) the closed face of EmrE does not appear to electrostatically coordinate the C

Further evidence of electrostatic gating is provided through investigation of the pH dependent behavior of EmrE when these gating residues are mutated. The rate of WT-EmrE alternating access is highly pH-dependent,^43,44^ with exchange slowing from the fast exchange limit (single peaks observed for each residue at the average chemical shift of the two exchanging conformations) with alternating access rate >200 s^-1^ at low pH (Fig. 4A, red) to the slow exchange limit (separate peaks observed at the unique chemical shift of each conformation) with alternating access rate <50 s^-1^ at high pH (Fig. 4A, blue). Along with increasing proton leak, truncating the C-terminal tail of EmrE (Δ107-EmrE) increases the rate of alternating access at high pH relative to WT-EmrE in the absence of substrate^14^. This increased rate of alternating access causes line broadening and merging of peaks in the 2D TROSY-HSQC spectrum of Δ107-EmrE at high pH (Fig. 4B, blue). This data shows that the C-terminal tail both gates proton and substrate access to the primary binding site at Glu-14 and regulates alternating access, suggesting that release of the tail from the fully occluded conformation is necessary for alternating access. If loop-tail electrostatic interactions are critical for maintaining the C-terminal tail gate in the occluded conformation, then mutations that disrupt these interactions should cause increased rates of alternating access relative to WT-EmrE and more closely resemble the Δ107-EmrE behavior. We recorded 2D ^1^H-^15^N TROSY-HSQC spectra of four conservative but neutralizing mutations of polar residues implicated in C-terminal tail gating by prior MD simulations or experimental studies of drug-substrate binding to the periphery of EmrE:^14,21,41,45^ E25Q-EmrE (TM1-TM2 loop), T56V-EmrE (TM2-TM3), D84N-EmrE (TM3-TM4 loop), and R106A-EmrE (C-terminal tail). Comparisons of these mutant spectra with WT-EmrE and Δ107-EmrE at both low and high pH are shown in Figure 4. E25Q-EmrE did not show any obvious change in the pH dependence of alternating access relative to WT-EmrE (Fig. 4C), but mutation of the other three other residues (T56V-EmrE, D84N-EmrE, and R106A-EmrE, Fig. 4D-F) did increase alternating access and cause similar spectra changes at high pH as observed in Δ107-EmrE. Thus, these three polar residues are vital to the C-terminal tail gating mechanism. This data is consistent with the Asp-84_B_ / Arg-106_A_ salt bridge and Thr-56_A_ / His-110_A_ hydrogen bond interactions in our structural model.

**Figure 4.**
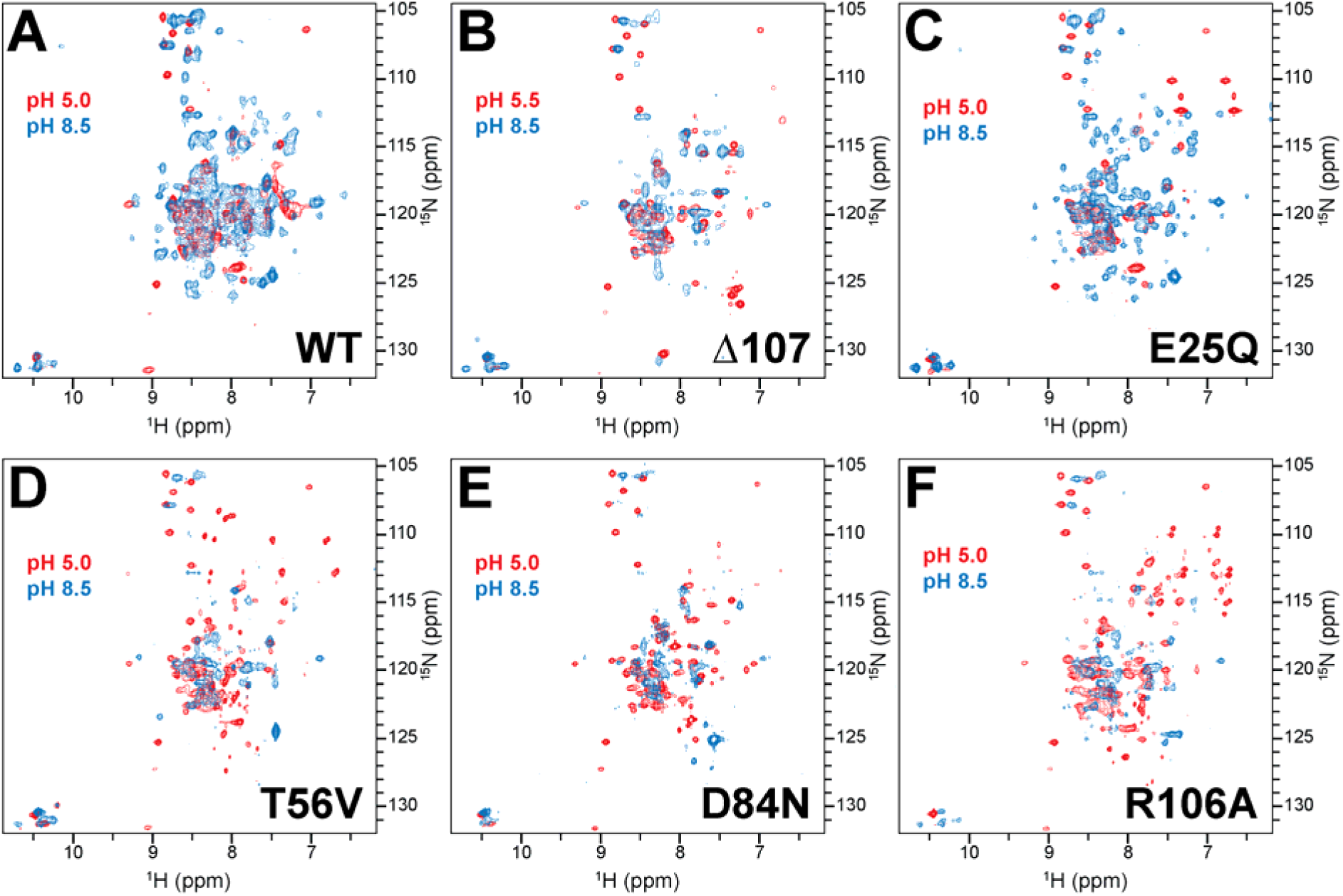
Electrostatic interactions stabilize tail conformation and regulate gating. At low pH (red spectra), all EmrE variants have 1 peak per residue, indicating fast (≥ 200 s^-1^) alternating access. At high pH (blue), WT-EmrE (A, blue) has 2 resolved peaks for most residues, indicating slow (≤ 50 s^-1^) alternating access. (B) The spectrum of Δ107- EmrE has more line-broadening at high pH (B, blue) than WT-EmrE (A, blue) and the distinct peaks for subunits A and B have coalesced to a single average peak for many residues, reflecting faster alternating access for Δ107-EmrE than WT-EmrE. The greater similarity between low- and high-pH spectra of Δ107-EmrE reflects less pH-dependent variation in alternating access rate for this mutant relative to WT-EmrE. (C) E25Q-EmrE has spectra similar to WT at high pH, suggesting that E25 has a less significant role in gating. T56V-EmrE (D), D84N-EmrE (E), and R106A-EmrE (F) all have substantial linebroadening and are shifted towards a single average peak for most residues at high pH, indicating faster alternating access than WT that is similar to Δ107-EmrE.

### The role of Ser-64 in regulating EmrE binding and dynamics

Alternating access in EmrE occurs when the two asymmetric subunits of the homodimer (chain A and chain B) swap conformations (AB to BA exchange). This conformational exchange is sufficiently fast to average peak positions in WT-EmrE at low pH (Fig. 4A, red) such that the unique peaks positions of the two subunits cannot be distinguished, thus precluding structure determination. At high pH, WT EmrE alternates access slow enough to resolve separate peaks for each subunit in the 2D ^1^H-^15^N TROSY spectrum (Fig. 4A, blue), but alternating access is still fast enough that line broadening degrades the ability to resolve all of the individual residues. Indeed, this significant exchange of the subunit conformations occurs during the mixing time of the NOESY experiments (solution NMR) or CC correlation experiments (solid-state NMR) such that cross-peaks arise from both conformers, resulting in distance restraints for each nucleus that are a mixture of both states. The S64V-EmrE variant slows the rate of alternating access rate of EmrE and reduces internal dynamics without disrupting substrate binding even though residue 64 is located near the middle of TM3 near the primary substrate binding site.^17,46^ This variant improves the quality of the NMR spectra, enabling chain and residue-specific distance restraint assignments without perturbing the loop and tail regions of EmrE. The significant shift in protein dynamics between WT- and S64V-EmrE observed with NMR motivated us to further characterize the functional role of the Ser-64 residue. Ser-64 is highly conserved in the QAC subfamily of SMR transporters such as EmrE^16,47^, but sequence alignment other SMR transporter subfamilies show that this residue can be glycine, alanine, threonine, valine, or even glutamate in other SMR homologs.^47,48^ To more fully characterize the impact of EmrE residue 64 on substrate interaction and transport, we mutated Ser-64 to Gly, Ala, Thr, or Val and measured ethidium efflux *E. coli*, *in vitro* TPP^+^ binding affinity, and *in vitro* rate of alternating access for the TPP^+^-bound transporter for each mutant (Table 2, Fig. 5, Fig. S4). All variants altered the transport activity in *E. coli* (Fig. 5A). Mutation to Gly, Ala or Val caused progressive slowing of ethidium efflux with increasing size of the amino acid side chain. Substitution of a very bulky tryptophan residue at this position is not tolerated and severely impairs the ability of EmrE to confer resistance to ethidium.^48^ Though bulkier, S64T-EmrE showed the most similar rate of efflux to WT-EmrE with just an initial delay. This indicates that the presence of the polar hydroxyl group on residue 64 is more essential for optimal efflux than a small side chain.

**Figure 5.**
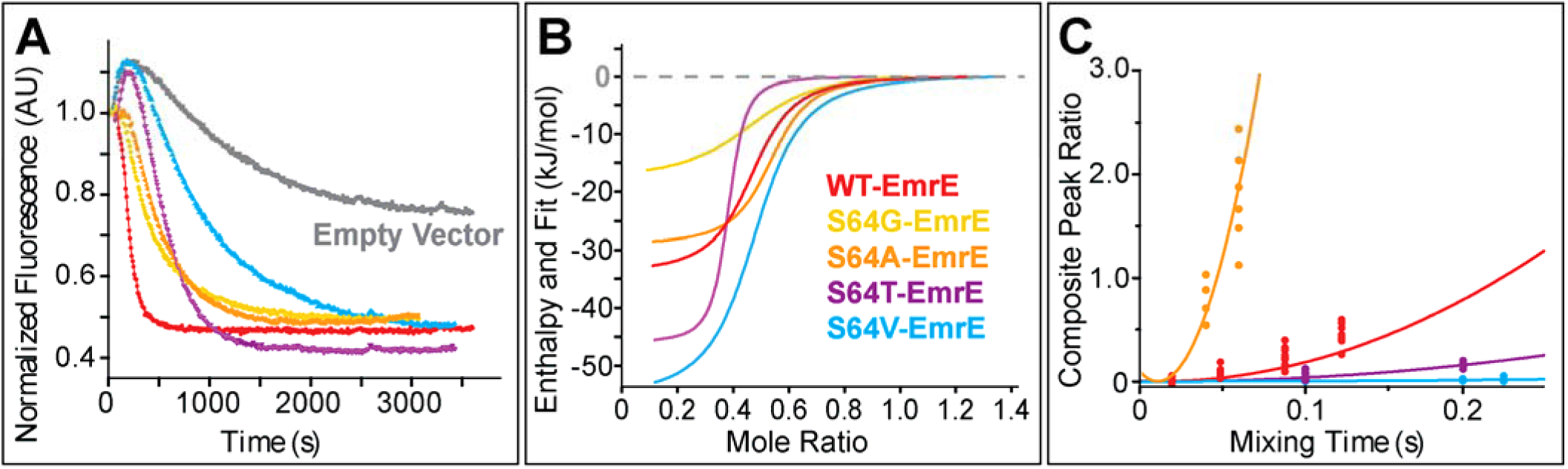
The residue at position 64 regulates EmrE substrate binding, alternating access and efflux. (A) Efflux of ethidium from *E. coli*. WT-EmrE (red) results in rapid efflux, while the empty vector control (gray) shows very limited efflux due to other multidrug efflux pumps exporting this common substrate. S64G-EmrE (yellow) and S64A-EmrE (orange) have the most similar initial rate of ethidium efflux to WT-EmrE. S64V-EmrE (blue) has much slower efflux but achieves a similar final internal ethidium concentration. S64T-EmrE (purple) has different initial behavior to WT-EmrE but then rapidly effluxes ethidium to a lower final baseline, demonstrating the importance of the sidechain hydroxyl group at residue 64 for efficient ethidium efflux. B) Representative ITC curves for EmrE binding to TPP+. Binding affinity is similar for all variants except S64G-EmrE (6-fold weaker affinity, Table 2) while binding enthalpy is similar for all variants except S64V-EmrE (modest 2-fold change, Table 2). C) ZZ-exchange NMR shows EmrE alternates access faster as the size of residue 64 decreases from S64V-EmrE (blue) to S64A-EmrE (orange) to S64G-EmrE (too fast to measure with this method). Full replicate data are shown in Figure S4.

**Table 2.** Impact of Ser-64 mutation on substrate binding and alternating access.

|  |  |  |  |
| --- | --- | --- | --- |
| **Residue 64** | **$K_d$ ($\mu\text{M}$)** | **$k_{conf}$ ($\text{s}^{-1}$)** | **$\Delta H_{\text{binding}}$ (kJ/mol)** |
| Gly | $1.5 \pm 0.1$ | $>150$ | $-17 \pm 2$ |
| Ala | $0.48 \pm 0.08$ | $21 \pm 5$ | $-30 \pm 2$ |
| Ser | $0.23 \pm 0.06^*$ | $4.5 \pm 0.5^*$ | $-22 \pm 1^*$ |
| Thr | $0.67 \pm 0.02$ | $2.4 \pm 0.2$ | $-27 \pm 4$ |
| Val | $0.51 \pm 0.07$ | $0.8 \pm 0.2$ | $-52 \pm 3$ |

\*Data from ref.<sup>56</sup>

Transport requires substrate binding, alternating access, and substrate release. We performed isothermal titration calorimetry (ITC) to investigate substrate binding and used NMR ZZ-exchange experiments to measure alternating access rates to determine whether the residue at position 64 had a greater impact on either of these factors. The ITC data (Fig. 5B, S4, Table 2) shows most of the Ser-64 mutations have little effect on substrate binding affinity. The one exception is S64G-EmrE, which decreases binding affinity by approximately 6-fold (Fig. 5B, Table 2). This could be due to the introduction of a third glycine (64-GGVG-67) in the middle to TM3, which may drastically increase helix flexibility disorder of the binding pocket. Binding enthalpy is similar across all variants, with the largest changes only 2- to 3-fold. These differences in binding affinity and enthalpy are small compared to the very large differences observed with minor modification of the substrate^49^ or mutation of residues that directly interact with substrate^40,50–53^ These results are consistent with the orientation of residue 64 away from the primary binding pocket (Fig. S5) while the neighboring residue, Trp-63, points directly into the binding pocket and is critical for substrate recognition and transport.^40,52,54,55^

More significant differences in alternating access rate are observed upon mutation of Ser-64 (Fig. 5C), supporting a role for this residue in regulating the conformational exchange between open-in and open-out states that is necessary for transmembrane transport. We used ZZ-exchange NMR to directly measure the rate of alternating access in EmrE as we have done previously.^12,17,42,49^ The data (Fig. 5C, Table 2) show a more than 100-fold increase in alternating access rate with decreasing amino acid side chain size from S64V-EmrE (0.8 s^-1^) to S64A-EmrE (21 s^-1^) to S64G-EmrE, (>150 s^-1^, too fast to measure with this method). S64A-EmrE also alternates access faster than WT, whereas bulkier S64V-EmrE and S64T-EmrE are noticeably slower than WT-EmrE. This collectively shows Ser-64 has both optimal size and polarity for EmrE drug binding and alternating access.

### The primary binding pocket of protonated S64V-EmrE is occluded from water

In all previous structures of EmrE determined using X-ray crystallography, electron microscopy and NMR,^22,23,25,27,32–34^ the primary binding pocket near Glu-14 in the core of EmrE appeared to be accessible to water on one side of the membrane, although the apparent degree of opening varied with the conditions under which the structure was determined, such as pH and refinement method. Such structures, with the binding pocket in the core of the transporter open and accessible to only one side, meet the requirement for active transport of a substrate against the electrochemical gradient.^4,5^ ^1^H-^15^N correlation spectroscopy has previously been used to quantify hydration of the EmrE binding pocket under different pH conditions,^23^ showing a significant increase in water accessibility into the core of EmrE at high pH. At low pH (Glu-14 protonated), those experiments indicated that the binding pocket was asymmetrically hydrated with only one side of the dimer accessible to water. These results were reinforced by MD simulations of substrate-free EmrE which estimated ∼47 water molecules in the binding cavity when both Glu-14 residues were deprotonated, compared to only ∼17 water molecules when both Glu-14 residues were protonated.^57^ These pH-dependent changes in hydration offer a mechanistic explanation for how EmrE regulates substrate access and proton exchange - low pH favors a dehydrated, more occluded state, while high pH promotes opening, water entry and release of protons. The structure presented here captures the dehydrated, gate-occluded state of EmrE found at low pH, and is the first structure of an occluded state for an SMR transporter.

To experimentally measure the water accessibility profile of substrate-free, protonated EmrE, we collected a series of ^13^C-^13^C SSNMR spectra and a T_2_ filter of 10 ms to eliminate polarization on protein and lipids and select for only water. By varying ^1^H-^1^H spin diffusion time to transfer polarization from water to EMRE via dipolar mediated interactions, the proximity to water may be measured as the build-up rate in a site-specific manner. Peak intensities were normalized to a control spectrum (no spin diffusion or T_2_ filter) and these relative peak intensities build-ups were fit to equation 4 (see Methods) to extract buildup rates with compensation for T_1_ in the fit (Fig. S7). Example build-up curves for individual residues are shown in Figure S6. As a complementary approach, we analyzed protein backbone amide-water NOEs from our 3D amide NOESY experiments. As the NOE depends on 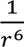 ^58^, data were plotted as log_6_ of the cross-peak intensities. These are plotted alongside the SSNMR buildup rates in Figure 6. This data demonstrates that the binding pocket of EmrE is highly dehydrated at low pH, consistent with previous computational studies of EmrE under these conditions,^57^ and provides further evidence of the C-terminal tail occluding access to the binding pocket of protonated EmrE.

**Figure 6.**
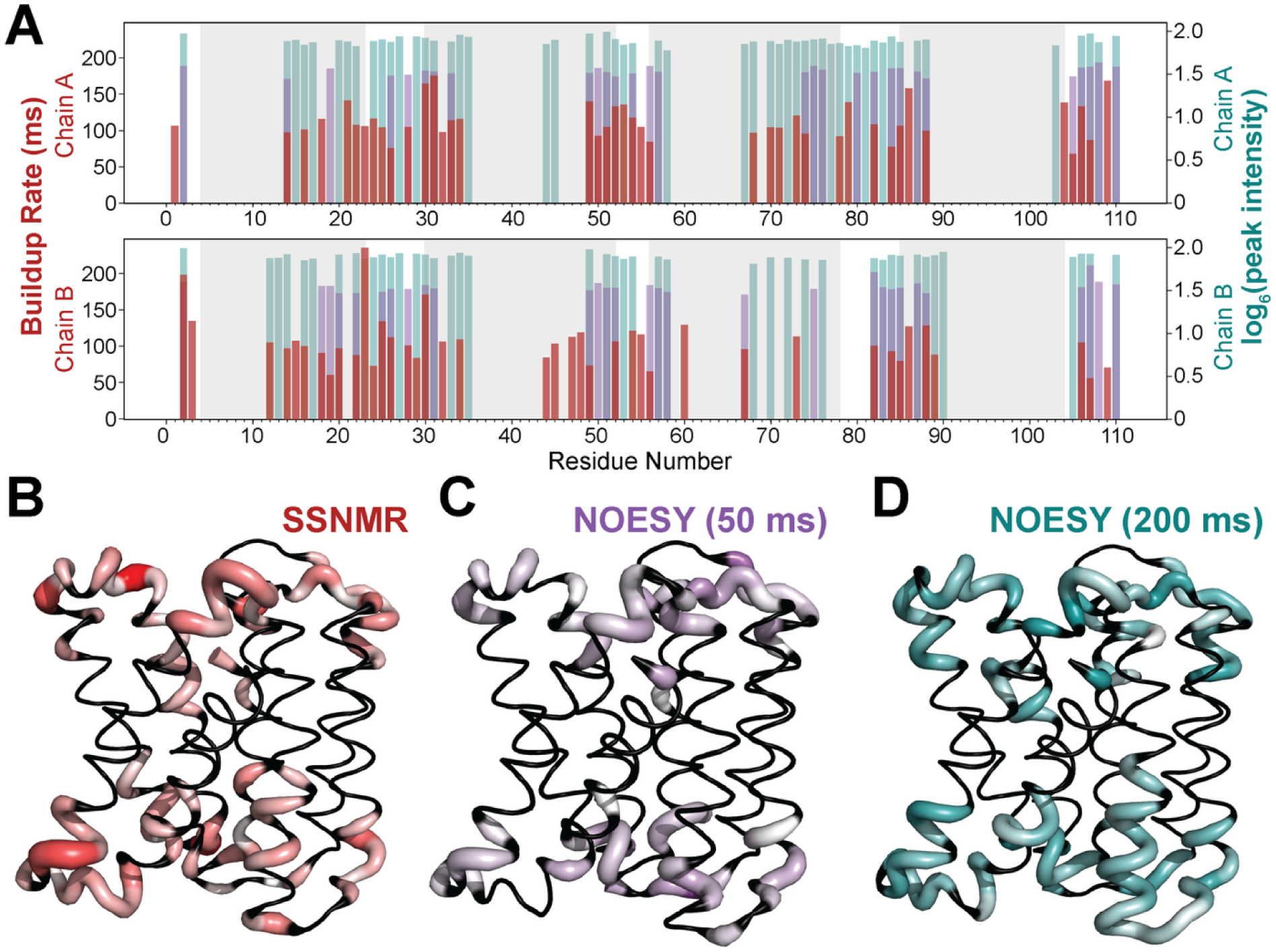
The protonated occluded state of EmrE has a mostly dehydrated binding pocket. (A) Water accessibility data is plotted per residue, including solid-state NMR buildup rates from spin diffusion experiments (ms, red), amide-water cross-peak intensities from the NOESY experiment with 50 ms mixing time (log6, purple), and amide-water cross-peak intensities from the NOESY experiment with 200 ms mixing time (log6, teal). The same data is plotted on the structure with scaled white to color and diameter proportional to magnitude of (B) SSNMR buildup rates, (C) 50 ms NOESY amide-water crosspeak intensity, and (D) 200 ms NOESY amide-water crosspeak intensity. The data reflects a lack of bulk water in the primary binding pocket near Glu-14 in the core of the transporter.

### Holistic evaluation of the available structural models of protonated EmrE

Our structural model of protonated S64V-EmrE confirms key features observed in two other experimental structures of protonated EmrE^22,25^ and adds the well-resolved structure of the C-terminal tail. This tail closes over the primary binding pocket, resulting in the first fully occluded structure of EmrE. The low pH crystal structure (PDB 7MH6) was solved at 2.58 Å using an EmrE variant with three mutations in the TM1-2 loop of EmrE and missing density for the C-terminal tail. At a strict 0 Å cutoff, this model satisfies 89% of our NMR-derived distance restraints, with excellent structural agreement within the TM helices (TM helix Cα RMSD 1.0 Å, Fig. S10). The NMR structure of protonated EmrE (PDB 8UWU) was determined using a heterodimer of WT-EmrE and L51I-EmrE, which has one mutation in the TM2-3 loop. This model also satisfies 91% ± 0.2% of our NMR-derived distance restraints, again primarily within the TM helices (TM helix Cα RMSD 2.2 Å). This model also had minimal restraints for the C-terminal tail but defined the structure and dynamics of key aromatic residues within the binding pocket of EmrE. Here we determined the structure of S64V-EmrE, which has fully native sequence for the loops and tails and a single mutation within TM helix 3. As noted above (Fig. 5, 7) and previously,^17^ this mutation does not significantly alter the structure of TM3 or the substrate affinity, it simply reduces the rate of alternating access. At a 0 Å violations cutoff, our low-pH, drug-free structure of EmrE satisfies 94% ± 0.6% of the measured distance restraints (Fig. 2, S2), which cover the full extent of EmrE including the loops and C-terminal tail. Thus, our structure is the most complete experimental model of protonated EmrE to date.

The C-terminal tail was previously shown to be functionally important in regulating proton leak, as deletion of only 4 amino acids at the end of the tail increased proton leak through EmrE to the extent that it impaired bacterial growth.^14,21,41^ Recent molecular dynamics simulations of EmrE identified electrostatic interactions between residues R106_A_ and D84_B_ as well as T56_A_, E25_B_ and H110_A_ that restrained the C-terminal tail in those models.^14,45^ Although the precise side chain conformations and importance of E25_B_ differs between the MD models and our structure, all of these models show the C-terminal tail interacts with loop residues to occlude water access to the primary binding pocket. Our work provides a structural basis for the observed role of the C-terminal tail in regulating proton flux and experimentally demonstrates that the C-terminal tail exists in a conformation that can gate access to the transport pore.

### Key structural features for EmrE alternating access and transport mechanisms

Previous cryo-EM and NMR studies have indicated that residues 65-GVG-67 within TM helix 3 of EmrE form a kink in one subunit of the homodimer^42,59,60^ that may act as a hinge point for alternating access. Mutagenesis of the two glycine residues in this region impair efflux and emphasize the importance of this region for enabling transport^17,40,48,50^ In agreement with these studies, our structure has a ∼30° kink at 65-GVG-67 of chain A (Fig. 7A). This kink is far less pronounced in chain B, where the angle at this junction is only ∼12°. The high resolution for both TM3 helices (1.5 Å ± 0.2 Å heavy atom RMSD in chain A and 1.1 Å ± 0.3 Å heavy atom RMSD in chain B) from extensive distance restraints from both solution and solid-state NMR data and backbone torsion angle restraints (BMRB accession numbers 52790 and 52793) reinforce to the fidelity of this kink. Other structural models of protonated EmrE also display a kink in TM3 at 65-GVG-67 in chain A, though to a lesser degree than our structure (Fig. S9).The kink in TM3 of chain A also fits well into the experimental crystallographic density of protonated EmrE,^22^ (Fig. 7B) demonstrating that the kink and adjacent regions are not significantly perturbed by the S64V mutation.

**Figure 7.**
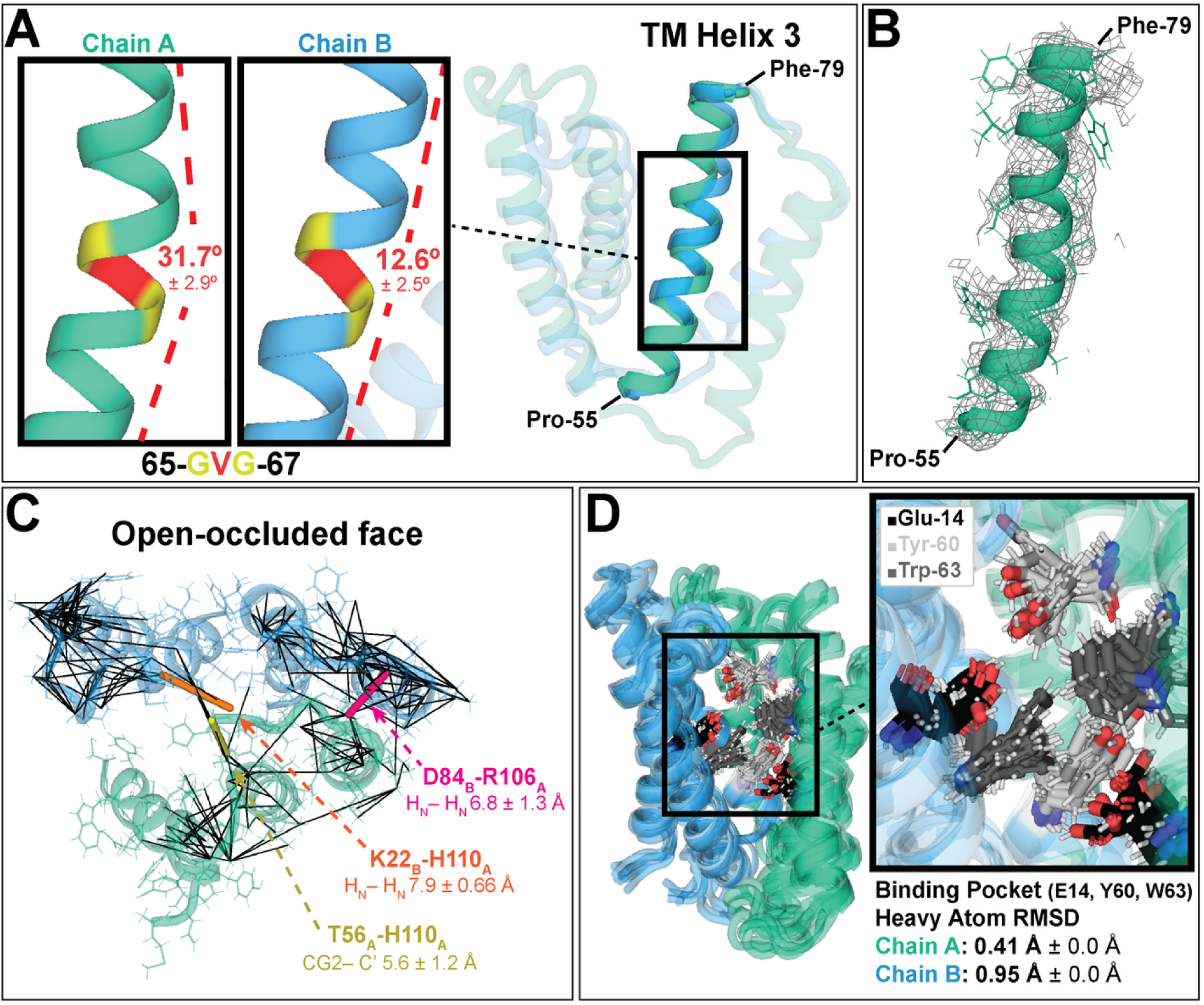
Functionally important structural features of protonated S64V-EmrE. A) A kink at 65-GVG-67 in TM helix 3 of chain A (not present in chain B) forms a key hinge point in the asymmetric homodimer important for alternating access of the primary binding pocket. B) The kink in our structure of S64V-EmrE at low pH (ribbon) fits well within previously published X-ray crystallography density of an EmrE variant with no mutation in TM3,1 confirming that the kink we observe is present in WTEmrE and the S64V mutation does not perturb EmrE structure in this important region. C) The C-terminal tail is held in place by electrostatic residues. Distance restraints near this gate are shown in black, and three key distance restraints between electrostatic residues are highlighted. D) Aromatic residues within the primary binding pocket are well aligned and tightly packed. The RMSD for specific binding pocket residues across the 10 lowest energy structures shows high resolution within the binding pocket of S64V-EmrE.

The S64V mutation slowing conformational exchange and its remoteness from the loops and C-terminal tail enabled structural determination of these functionally important regions of EmrE in the absence of mutations or potentially structurally perturbing antibodies to this region. Solution and solid-state NMR data revealed site-specific distance restraints in these regions, including between loop and tail residues. For example, in the 200ms mixing time amide ^15^N-^1^H-^1^H NOESY NMR spectra we observed clear NOE crosspeaks between the backbone amide protons of Asp-84_B_ and Arg-106_A_ and of Lys-22_B_ and His-110_A_, and in the 3D ^13^C DARR experiments, we observed multiple carbon correlations between Thr-56_A_ and His-110_A_ (Fig. S9). This allowed us to directly restrain loop and tail residues with functional importance (Fig. 7C).

Although the C-terminal tail is not present or not well-resolved in previous experimental structures of EmrE, a recent NMR structural model^25^ found the binding pocket of protonated EmrE to be partially occluded to drug-substrate, with aromatic residues Tyr-60_A,B_ and Trp-63_A,B_ packing tightly against Glu-14_A,B_ to block substrate access. This is consistent with our water accessibility experiments showing a dehydrated binding pocket, and with the unusual highly-elevated pKa values of Glu-14_A,B_^61^ since these aromatic residues create a highly hydrophobic local environment. Similarly, our low pH occluded structure of EmrE also has tight packing of aromatic residues Tyr-60_A,B_ and Trp-63_A,B_ with the canonical binding residues Glu-14_A,B_ (Fig. 7D) in 9 out of the 10 lowest energy structures. In only one of our ten lowest energy structures do we see a Trp-63 indole ring flip, supporting the previous conclusion that equilibrium largely favors an occluded binding pocket.^25^

## CONCLUSION

Structures of secondary active transporters in different states are necessary for developing three-dimensional understanding of coupled transport.^5^ The structural and biophysical data reported in this study revealed that the C-terminal tail gates proton leak through substrate-free EmrE via electrostatic interactions, and we showed that our S64V point mutation does not perturb the structure of EmrE. The structure of protonated EmrE presented here is the first occluded experimental structure of EmrE and of an SMR transporter and supports previous suggestions that SMR transporters must have an occluded state to prevent uncoupled proton leak during the process of alternating access.^14,62,63^ This work further highlights how complex these apparently “simple” model transporters are, and that sophisticated gating can be achieved even within a tiny transporter.

## MATERIALS & METHODS

### EmrE expression and purification

All EmrE samples for solution- and solid-state NMR were expressed, purified, and reconstituted exactly as described in the assignment note corresponding to this manuscript.^46^ EmrE samples for S64 mutant functional assays were expressed and purified in the same fashion and as previously described.^17,64^ Briefly, a His-tagged EmrE pet15b construct with the appropriate mutation was expressed in BL21(DE3) *E. coli*, grown in M9-minimal media with appropriate isotopic enrichment, then induced overnight at 17°C with 330 µM isopropyl beta-D-thiogalactopyranoside (IPTG). EmrE was purified in n-decyl-β-maltoside (DM) with nickel affinity chromatography, His-tag was cleaved with thrombin (Sigma), and EmrE was further purified with size exclusion chromatography (SEC) as previously described.^64^ Fractions containing purified EmrE were sonicated with long chain lipid and octylglucopyranoside (OG) to form proteoliposomes and detergent was removed with Amberlite (Supelco) overnight. Proteoliposomes were collected via ultracentrifugation. Dimyristoylphosphatidylcholine (DHPC) was added to samples for solution NMR to form isotropic bicelles (q=0.33). All samples were subjected to a freeze-thaw cycle in liquid N_2_ at least three times before use.

### Solution NMR restraints

Solution NMR experiments were collected and analyzed exactly as described in the assignment note corresponding to this manuscript.^46^ Two further 3D ^1^H, ^1^H amide NOESY experiments were collected with 25% non-uniform sampling (NUS)^65,66^ on a ^2^H/^15^N S64V EmrE samples. One was collected using a Bruker NOE spectrometer at 1.1 GHz equipped with a 3 mm TCI cryoprobe with the following parameters: ns=32, d1=1.5s, d8=200ms. The other was collected on a Bruker Avance III HD spectrometer at 900 MHz equipped with a 5 mm TCI cryoprobe with the following parameters: ns=16, d1=2s, d8=50ms. Both amide NOESY spectra were processed with NMRpipe^67^ using SMILE reconstruction^68^ and analyzed with CCPNMR v3.0.^69^

Distance restraints were extracted by matching crosspeaks in the 3D amide, 3D methyl, and 4D methyl NOESY experiments. Peaks in the 3D amide NOESY experiment often overlapped, so atoms with crosspeaks were given a blanket 2-8 Å distance range. Peaks in the 3D methyl NOESY experiments often overlapped, so atoms with crosspeaks were given a blanket 2-12 Å distance range. Peaks in the 4D methyl NOESY experiment were largely highly resolved, so distance restraints were calibrated to peak volume. Distance restraints were given a “starting point” of 6 Å then individual peak volumes were normalized to the median peak volume and converted to distance values in Å according to Equation 1:

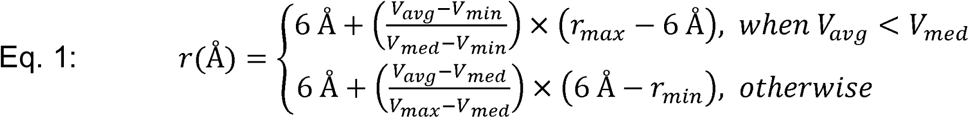

where V_avg_, V_min_, V_med_, and V_max_ correspond to peak volume average, minimum, median, and maximum respectively, and r_max_ and r_min_ correspond to user-defined maximum and minimum distance values (in Å) respectively. Calculated distance values were given ± 2Å of flexibility and input as distance restraints in structure calculations with XPLOR-NIH.

### Solid-State NMR restraints

Solid-state NMR (SSNMR) experiments were collected and analyzed exactly as described in the assignment note corresponding to this manuscript.^46^ A further 3D NCOCX (600 ms, DARR) was collected on a 17.6 T (750 MHz ^1^H frequency) wide bore magnet equipped with a Varian Balun probe in HCN mode and spun at an MAS rate of 12.5 kHz at a sample temperature of -23 ± 5 C. The ^1^H, ^13^C, and ^15^N pulse widths were 2.2 us, 2.3 us, and 4.2 us, respectively. The ^1^H to ^15^N CP field strengths were 51.4 kHz and 52.5 kHz, respectively, with a downward tangential ramp on the ^1^H channel (Δ = -10.5 kHz and β = 2.3 kHz) with a contact time of 1.8 ms and the ^15^N carrier frequency set to 98.03 ppm and a sweep width of 6250 Hz and digitized out to 16 ms. The ^15^N to ^13^C CP field strengths were 20.5 and 42.1 kHz, respectively with an upward ramp on the ^15^N channel (Δ = 1.8 kHz and β = 1.3 kHz) with a contact time of 6 ms and the C’ frequency offset set to 172.3 ppm and a sweep width of 6250 Hz and digitized out to 12.8 ms. Homonuclear recoupling of 600ms using DARR was used to identify inter-residue contacts and the directly detected dimension was acquired for 18 ms with a sweep width of 100 kHz. SPINAL-64 decoupling (*ω*_1_ ^H^ = 82.0 kHz) was employed during data acquisition. A 50% Poisson gap (2x sine bell weighted) schedule was generated using nus-tool and employed to collect approximately six replicates separated by 1D HNCO reference spectra to ensure CP stability (∼38 hr each block). The spectrum was processed using NMRPipe^67^ using sine bell (90° offset) window functions in all dimensions and reconstructed using hmsIST.

### Transferred Echo Double Resonance (TEDOR)

A series of 2D ^15^N-^13^C planes were acquired according to the 3D ZF-TEDOR pulse sequence.^70^ SPINAL-64 decoupling was used during acquisition with a ^1^H field of 82.4 kHz (5.5 us with a 5° phase increment), and TPPM decoupling was employed during REDOR (5.0 us, 8.979°) at a ^1^H field of 90.9 kHz. ^13^C and ^15^N π pulse widths during TEDOR were 4.6 us and 15 us respectively. Spectra were acquired with 16 scans per row, resulting in a 3.5 hr block with each 2D plane digitized to 512 points (TPPI) by 80 us in t1 (15N, 20.48 ms maximum evolution time) and 1800 complex points (10 us dwell) in the t2 acquisition dimension (^13^C, 18 ms acquisition time). The ^15^N-^13^C dipolar trajectories were sampled with values of tmix = 1.28. 2.56, 3.84, 5.12, 7.68, 8.96, and 10.12 ms. Each mixing time was collected for at least 12 blocks (3.5 hr each, 42 hours total).

Data were processed in NMRPipe using 54° sine bell offsets in each dimension and the indirect dimension was extended out to 30.7 ms (384 points) using SMILE.^68^ Spectra were assigned n NMRFAM-SPARKY^71^ and the peak heights at the same position in each spectrum were measured and fit the following TEDOR buildup curve derived from Jaroniec et al.^72^ (Eq. 2):

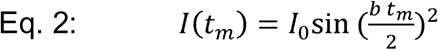

where *I_0_* is the scaling factor, *b* is the dipolar coupling constant, and *t_m_* is the TEDOR mixing time. Only crosspeaks with an R² value greater than 0.8 were included in the final analysis. Nonlinear least-squares curve fitting was performed with initial parameter guesses based on the maximum observed intensity and estimated oscillation periods. Dipolar couplings obtained from the fits were converted to distances using the point-dipole approximation. Distances were calculated according to Equation 3:

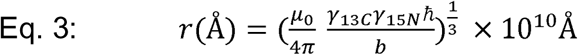

where γ13*C* and γ15*N* are the gyromagnetic ratios of ^13^C and ^15^N, respectively, ℏ is the reduced Plank’s constant, and 𝜇 is the permeability of free space. For each assignment, the fitting results and corresponding TEDOR distances were recorded, and the fitted curves were plotted alongside the experimental data (Fig. S6). Distances were then converted to Xplor-NIH format and used for distance restraints in the calculations.

### SSNMR Water Accessibility Experiments

Uniformly labeled ^13^C,^15^N EmrE in lipid bilayers were packed in a 2.5 mm rotor and ^13^C-^13^C spectra with various spin diffusion times were collected at 900 MHz using a 2.5 mm BlackFox probe at an MAS rate of 20 kHz. The variable temperature (VT) was set to -10 °C resulting in an effective sample temperature of -3 °C. ^1^H and ^13^C 90° pulse widths were 3.75 and 3.4 us, respectively. Spectra were collected with 1.2 ms CP contact time with nutation frequencies 𝜈C = 90 kHz and a downward tangent ramp on ^1^H (Δ = -1.5 kHz and β = 0.8 kHz) with an average RF amplitude of 85 kHz. There is a 10% error when extracting tangential shape parameters Δ and β. Homonuclear recoupling used CORDxy4, 100 ms.^73^. The maximum evolution time (t_1_) was 3.2 ms (6.3 µs x 512 total points), and the t_2_ acquisition time was 20.5 ms (5 µs x 4096 total points). Spectra were acquired with a ^1^H T2 filter of 10 ms with spin diffusion times of 16, 36, 64, 100, 144, 256, and 400 ms. The recycle delay was 1.5 s with 32 scans per row and repeated four times. The total experimental time was 32 hours per series of spectra collected with a single spin diffusion time. All experiments utilized 90 kHz SPINAL-64 ^1^H decoupling during evolution and acquisition times.

Water accessibility data was processed by assigning peaks in the ^13^C-^13^C (100 ms) spectrum with 144 ms of spin diffusion which contained the most peaks. The peak intensities at the same position in the other ^13^C-^13^C water accessibility spectra were also measured and fit to a simplified equation defined by Amani et al.^74^ (Equation 4):

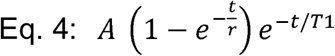

where *A* is the scaling factor fit to the experimental data, *t* is the spin diffusion time (s), *r* is the buildup rate (s^-1^) fit to the experimental data, and T1 are the T1 values measured for methyl groups (800 ms), carbonyl groups (782 ms), and other backbone and sidechain groups (measured from 40 to 75 ppm, 796 ms). T1 inversion recovery data is included in Figure S7. For each assigned peak, the buildup rate was applied to the ^13^C atom in the direct dimension, followed by calculation of the average buildup rate for each residue. The average buildup rate per reside was converted to Xplor-NIH format and implemented into the Xplor potential PSolPot.^31^

### Automated assignment of 3D SSNMR spectra with PASD

We employed the probabilistic assignment algorithm for automated structure determination (PASD) algorithm^75,76^ as a tool to assign our 3D SSNMR experiments including ^13^C-^13^C-^13^C (DARR, t_mix,1_ = 50 ms and t_mix,2_ = 50 ms) and ^13^C-^13^C-^13^C (DARR, t_mix,1_ = 50 ms and t_mix,2_ = 250 ms) experiments performed on uniformly ^13^C labeled S64V-EmrE, a ^13^C-^13^C-^13^C (DARR, t_mix,1_ = 150 ms and t_mix,2_ = 600 ms) experiment performed on ^13^C 1,3-glycerol labeled EmrE above the phase transition temperature of POPC, and a NCOCX (DARR, t_mix_ = 600 ms) experiment performed on ^13^C 1,3-glycerol labeled EmrE below the phase transition temperature of POPC. PASD was performed in accordance with previous studies.^77^ Preliminary structure calculations were first performed using the Xplor-NIH software package (version 3.6.6) employing a simulated annealing protocol. Initial coordinates were derived from a modified homology model based on PDB entry 8UWU with mutations reset to the WT sequence then addition of mutation S64V. Calculations incorporated a potential energy function including standard covalent geometry (bonds, angles, impropers), torsion angle database potentials, and EEFx implicit membrane solvation potentials^78^ (param_LK36). Dihedral angle restraints were derived from solution TALOS-N predictions, and distance restraints were obtained from solution NMR spectra. A membrane environment was modeled using IMMx settings with a 27 Å membrane thickness and default dielectric screening parameters. A multi-stage simulated annealing procedure was implemented, starting with high-temperature dynamics using coarse van der Waals (VDW) repulsion (onlyCA=1), followed by gradual cooling and ramping up of full-atom potentials, including VDW, and dihedral restraints. One hundred structures were calculated with random seeds assigned based on process ID for stochastic sampling.

We then executed the PASD protocol in Xplor-NIH, using the lowest energy structure from the solution NMR-driven structure calculations as the starting point. Cross-peak intensities were matched to assigned resonance frequencies using 0.2 ppm tolerances in both direct and indirect dimensions. Automated chemical shifts correction was not performed. Binned distances used for PASD are shown in Table S1. A network filter was applied to assign likelihoods of either 0 (not assigned) or 1 (assigned) to each peak. This filter reduced restraint ambiguity and emphasized high-confidence, spatially localized NOE assignments, thereby improving the reliability of downstream structural models. For peaks with at least one intraresidue assignment, all non-intraresidue assignments were discarded to prioritize structurally local interactions. The filtered peaks, exceptions, and chemical shift assignment files were saved for subsequent PASD structure calculations.

PASD calculations were performed by employing a non-crystallographic symmetry (NCS) position differential potential on the helices to keep them held in position relative to one another while randomizing the coordinates of the interhelical loops and C-terminal tail. Calculations of 500 structures for each pass were performed, and statistics were collected on the 200 lowest energy structures for each pass. Peak assignment likelihood reanalysis following each PASD structure calculation pass was done using the 20 best-converged structures from the low energy fraction. Pass 2 and pass 3 were run using only peaks with an atom-atom peak assignment ambiguity of 1 or 2. Likelihood reanalysis based on the pass 3 structures was done using the pass 1 peak lists, which re-introduced the originally higher ambiguity peaks, now at lower ambiguity. High-likelihood restraints (≥ 0.9) were kept. Restraints which varied >0.5 ppm from their resonance list assignments were discarded, and remaining long range inter-residue (|i – j| ≥ 5) restraints were manually verified with SSNMR spectra. Final restraints were converted to Xplor-NIH format for structure calculations.

### Xplor-NIH structure calculations

The final simulated annealing protocol began with an extended structure which was folded using NMR restraints and previously published X-ray crystallography density of low-pH substrate-free EmrE^22^ as potentials. The following NMR restraints were included as potentials for structure calculation with Xplor-NIH: solution NMR NOESY distance (3D amide NOESY, 3D methyl NOESY, and 4D methyl NOESY), SSNMR restraints derived from PASD, per-residue water accessibility derived from SSNMR, and internuclear ^13^C-^15^N distance restraints from TEDOR. A paramagnetic solvent accessibility potential (PSolPot) was used to refine the structure against experimental water accessibility data. The potential was created using create_PSolPot from the Xplor-NIH psolPotTools module, with input from a normalized solvent accessibility table. A probe radius of 4.0 Å and a fixed correlation time (τc) of 0.2 ns were used to model the effects of a paramagnetic solvent. The potential was evaluated using a correlation-based scoring function, with additional parameters set to mimic Gd³⁺-like relaxation behavior (S = 3.5, ρ₀ = 4). The PSolPot energy term was included in the total energy with a weighting scale of 10. A pseudo-energy potential combining covalent geometry terms (bond, angle, improper), torsion angle preferences from a database potential, and TALOS-N dihedral restraints were employed. Nonbonded interactions were modeled using a van der Waals repulsion potential with a staged transition from CA-only to all-atom resolution. The annealing schedule consisted of three high-temperature stages followed by a gradual ramping of potential energy terms, including VDW, torsionDB, and repel potentials, to their final target values. One hundred structure calculations were run in total. The ten structures with the lowest energy potential showed an average heavy-atom pairwise RMSD of 2.13 Å.

### Structure minimization and equilibration with GROMACS

The 10 Xplor-NIH output structures with the lowest overall energy potential were input into explicit POPC lipid using CHARMM-GUI^36^ Bilayer Builder server for minimization and equilibration with GROMACS.^37^ PDB files were uploaded then aligned in the bilayer using PPM 2.0.^79^ pH was set to 5.8, and residues with sidechains known to be protonated (Glu-14_A_ and Glu-14_B_^44^, His-110_A_ and His-110_B_^16^) were protonated. Pore water was generated using protein geometry and extraneous water molecules were removed manually. A rectangular lipid box was constructed with roughly 1/3 protein surface area and 2/3 lipid surface area (X and Y = ∼65). The system was solvated, and neutralizing sodium ions were added at 0.2 M concentration. Temperature was set to 318.15 K. NPT ensemble output files for GROMACS were downloaded for each of the ten structures. Software xplor2gmx from the Debsources Dataset^80^ was used to convert Xplor-formatted distance restraints to GROMACS position restraint (posre) potentials, and in-house scripts were used to convert TALOS-derived dihedral angles to GROMACS dihedral restraints then given a force constant of 1000 kj/mol•nm^2^. Each structure was minimized using the steepest decent algorithm and equilibrated for 875 ps in the presence of NMR distance restraints and dihedral restraints. Coordinates for each EmrE structure on the last frame of equilibration were extracted and combined into our final structural model bundle.

### Structure evaluation

Distance restraint and torsion angle violations were evaluated using VMD-XPLOR^81^ with a cutoff of 0 Å and 0° respectively. Deviations from idealized geometry were evaluated with PDBstat.^82^ Average pairwise RMSD was determined with MDAnalysis.^83–86^ Hydrogen bonds and salt bridges were evaluated with VMD-XPLOR.^81^

### Loop and C-terminal tail mutant NMR

For each loop/tail mutant, two-dimensional amide TROSY^87^ NMR spectra were collected on 0.5-1.0 mM uniformly labeled ^2^H^15^N EmrE in DMPC/DHPC bicelles (q = 0.33, with a protein to DMPC molar ratio of 1:75) in buffer consisting of 20 mM NaCl, 20 mM acetate, 50 mM MOPS, and 50 mM bicine. Samples were split and prepared at pH 5 and pH 8.5, then NMR experiments were acquired. To verify sample stability, the two samples were mixed to create intermediate pH values, and NMR data was acquired across intermediate pH values (data not shown).

### Chemical Shift Perturbation (CSP) analysis

CSP analysis between chains A and B of S64V-EmrE was conducted as previously described.^56^ Differences in ^1^H chemical shifts (Δδ*_HN_*) and ^15^N chemical shifts (Δδ*_N_*) between chain A and chain B of EmrE were weighted according to the standard formula^60,88^ shown in Equation 5:

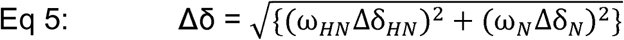

where ω_i_ is the weighting factor for each nucleus (ω_HN_ = 1, ω_N_ = 0.154) and Δδ is the CSP between chain A and chain B. A significance cutoff of 1 SD from the mean Δδ value for all residues was used to differentiate significant and insignificant chemical shifts. For reference, this distribution is plotted in Figure S11. Unassigned residues were excluded from analysis.

### S64 mutant functional assays

In-cell ethidium efflux assays were carried out using BL21(DE3) *E. coli* transformed with empty pET15b vector or pET15b containing WT or S64 mutant EmrE. Cells were grown in M9 minimal media with 100 μg/ml of ampicillin at 37 °C until the OD_600_ reached 0.4. Cells were induced with 0.33 mM isopropyl 1-thio-β-D-galactopyranoside (IPTG) for 30 minutes at 37°C. 2.5 μM ethidium bromide and 40 μM carbonyl cyanide p-chlorophenylhydrazone (CCCP) were added and the cells were incubated at 37°C for an additional hour. Assays were started immediately with excess cell culture stored on ice until it was in an assay. For each experiment, 2 mL of cell culture was spun down and immediately resuspended in 1 ml fresh M9 media with 2.5 μM ethidium bromide. Ethidium bromide fluorescence was monitored with excitation at 545 nm and emission at 610 nm. Fluorescence values over time were normalized to the initial fluorescence value of each run.

For each S64 mutant, ZZ-exchange NMR data was collected on 0.8-1.5 mM uniformly labeled ^2^H^15^N EmrE in DMPC/DHPC bicelles (q = 0.33, with a protein to DMPC molar ratio of 1:75) bound to saturating 2 mM TPP^+^. Buffer conditions consisted of 100 mM MOPS, 10-30 mM NaCl, 2 mM TCEP, 8-10 % D_2_O, pH 7 at 45 °C. Experiments were collected at 45°C on a Varian 700 MHz spectrometer with a room temperature probe. All NMR spectra were processed with NMRPipe^67^ and analyzed in CcpNmr Analysis.^89^ 2D ^1^H,^15^N TROSY-HSQC and TROSY-selected ZZ-exchange experiments^90^ with a lipid flip-back pulse^91^ were carried out with a recycle delay of 2 s and 128-144 increments. The conformational interconversion rate, *k_conf_*, was determined as previously described^92^ using the composite peak ratio method with an 11.1 ms offset time, *t_0_*, to account for the back-transfer time in the pulse sequence.^93^ The composite peak ratios of intensities of the auto-peaks (*I_AA_*, *I_BB_*) and cross-peaks (*I_AB_*, *I_BA_*) were fit to Equation 6 as a function of the delay time, *t*. Two planes with different mixing times were collected for each mutant, and mixing times were adjusted according to the conformational interconversion rate of each mutant. The mixing times used were 40, 60 ms for S64A, 100, 200 ms for S64T, and 200, 225 ms for S64V.

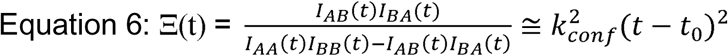

Isothermal titration calorimetry (ITC) experiments were run at 45 °C with a TA Instruments Low Volume Nano ITC calorimeter and operated with ITCRun software (TA Instruments) as described previously ^42,56^. Briefly, the reference cell was loaded with 350 µL degassed MQ H_2_O and the sample cell was loaded with 300 µL 1:3 1,2-dimyristoyl-*sn*-glycero-3-phosphocholine (DMPC): dihexanoylphosphatidylcholine (DHPC) bicelle-reconstituted WT, S64G, S64A, S64V, or S64T EmrE at monomer concentrations of 20-50 µM. The stirring rate was set to 350 rpm. The titrant syringe was loaded with 50 µL of ∼100-400 µM TPP^+^ mixed with bicelles which matched the composition and concentration of bicelles used in the corresponding EmrE sample. Titrant was injected in 2.5 µL aliquots every 5 minutes for a total of 20 injections. This ensured a constant lipid and buffer condition to minimize any heat of mixing or partitioning of substrate into lipids. Baseline correction, fitting, and data analysis were conducted using NanoAnalyze software (TA Instruments). Three replicate experiments were collected and analyzed per mutant. Raw heats were baseline corrected and then fit to Equation 7 to determine the association constant (*K*) and dissociation constant (*K_d_* = *K*^-1^), enthalpy of binding (ΔH), and binding stoichiometry (*n*).

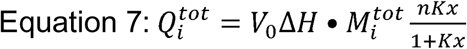

where Q^i^_tot_ is total heat after the i^th^ injection, V_0_ is cell volume, M^i^_tot_ is total EmrE monomer concentration after the i^th^ injection, and x is ligand TPP^+^ concentration.

## ACKNOWLEDGEMENTS

The authors thank M. Tonelli and S. Wang for assistance with NMR data collection, P. Liu for assistance with XPLOR-NIH and M. O’Connor for assistance with GROMACS. Sample preparation and data acquisition was supported by National Institute of General Medical Sciences of the National Institutes of Health with award number R35GM141748 to KHW. This work utilized instrumentation from the National Magnetic Resonance Facility at Madison, which is supported by National Institutes of Health grants R24GM141526 and P41GM136463. Data acquired at 1.1 GHz NMR and infrastructure for NAN data archive used equipment funded by the United States National Science Foundation Mid-Scale Research Infrastructure RI-2 program under Grant No. 1946970. Data acquired at 900 MHz used equipment funded by the National Institutes of Health S10OD034243. ABHB was supported by Molecular Biophysics Predoctoral Training Grant T32GM130550 from the National Institute of General Medical Sciences and the Dr. James Chieh-Hsia Mao Wisconsin Distinguished Graduate Fellowship administered by the University of Wisconsin-Madison Department of Biochemistry. BDH was supported by the Molecular Biophysics Predoctoral Training Grant T32GM130550 from the National Institute of General Medical Sciences and the William H. Peterson Graduate Fellowship and Steenbock Predoctoral Graduate Fellowship administered by the University of Wisconsin-Madison Department of Biochemistry.

## Contributions

ABHB prepared EmrE samples, performed and analyzed solution-state NMR experiments, completed manual verification of NMR data and structure calculations with XPLOR-NIH and GROMACS, analyzed ZZ-exchange NMR and ITC data, critically evaluated the structural model and drafted the manuscript; BDH performed and analyzed solid-state NMR experiments, initialized XPLOR-NIH structure calculation scripts, and assisted with drafting the manuscript. MB performed solution NMR pH titrations. OW assisted with XPLOR-NIH script development. CGB assisted with GROMACS. CW performed ZZ-exchange NMR. EU collected ITC data. CCC performed short-mixing time amide NOESY data. GR assisted with EmrE-mutant experiments. CMR collected solid-state NMR data and provided funding and oversight. KAHW provided funding, oversaw data analysis and interpretation and revised the manuscript.

**ACCESSION CODES**

**Primary Accessions**

**Protein Data Bank (PDB)** 36AV

- DOI: 10.2210/pdb36av/pdb

**BMRB**

- Solution NMR: 52790
- Solid-state NMR: 52793

**BMRBig**

- Solution NMR: bmrbig109
- Solid-state NMR: bmrbig110

**Network for Advanced NMR (NAN)**

- Solution NMR: usnan.org/ark:/83454/c1c5690d6c-b839-4b37-82a4-393e903501e3
- NCACO: usnan.org/ark:/83454/c19bbc0d40-3f21-453e-bb11-39452d726f03

**Referenced Accessions**

**Protein Data Bank** 7MH6, 7MGX, 7SVX, 7SSU, 7SV9, 7T00, 7JK8, 7SFQ, 8UWU, 8OUZ

