## Supplementary Material for "Protonated Structure of EmrE Reveals C-terminal Tail Gating Mechanism"

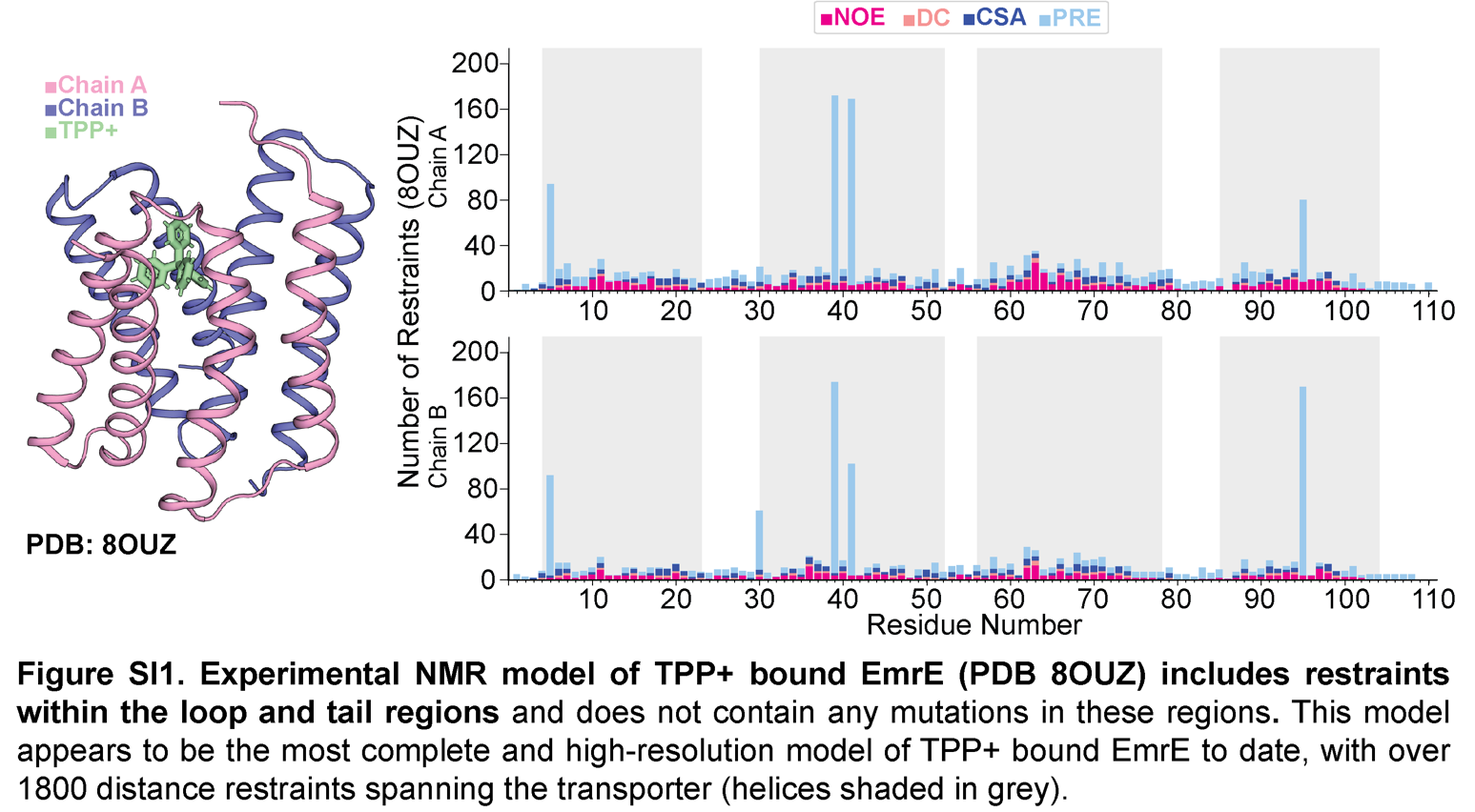

**Figure S1. Experimental NMR model of TPP+ bound EmrE (PDB 8OUZ) includes restraints within the TM loop regions of EmrE** and does not contain any mutations in these regions (helices shaded in grey).

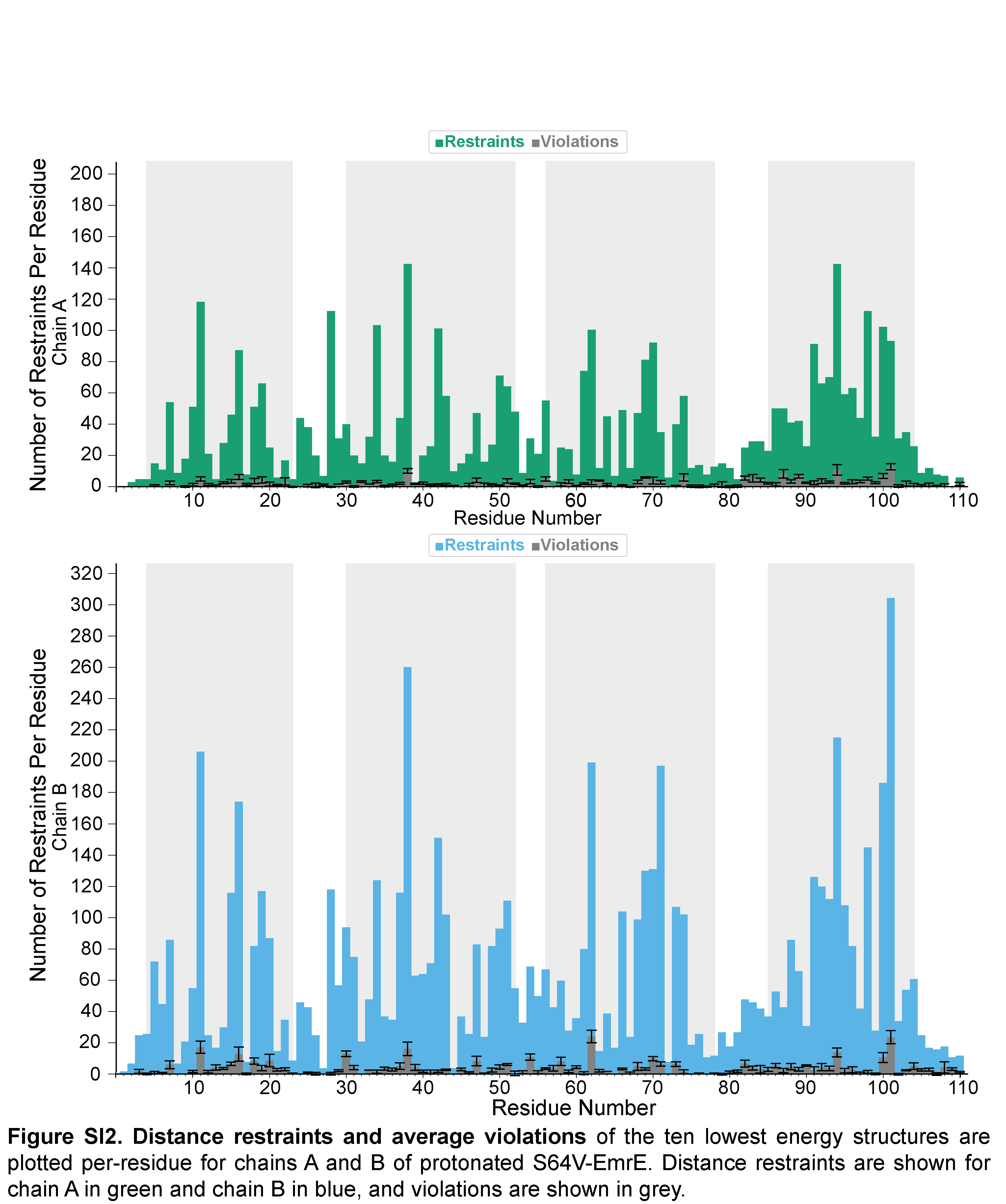

**Figure S2. Distance restraints and average violations of the ten lowest energy structures** are plotted per-residue for chains A and B of protonated S64V-EmrE. Distance restraints are shown for chain A in green and chain B in blue, and violations are shown in dark grey. TM helices are shaded in light grey.

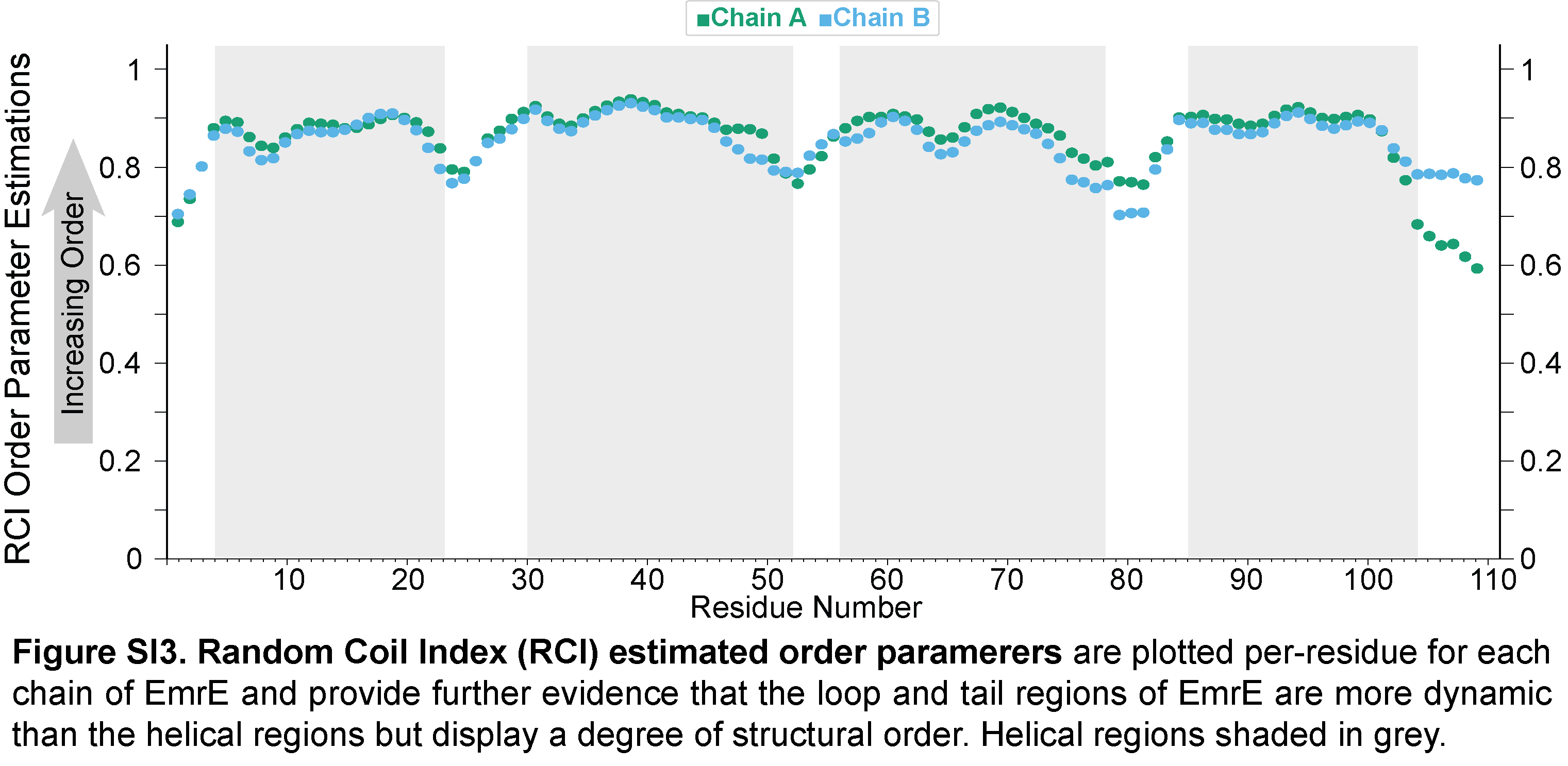

**Figure S3. The loop and tail regions of EmrE are dynamic but not disordered.** Random Coil Index (RCI) estimated order parameters are plotted per-residue for each chain of EmrE (1 = rigid; 0 = fully disordered) and provide further evidence that the loop and tail regions of EmrE are more dynamic than the helical regions but display a degree of structural order. Helical regions shaded in grey.

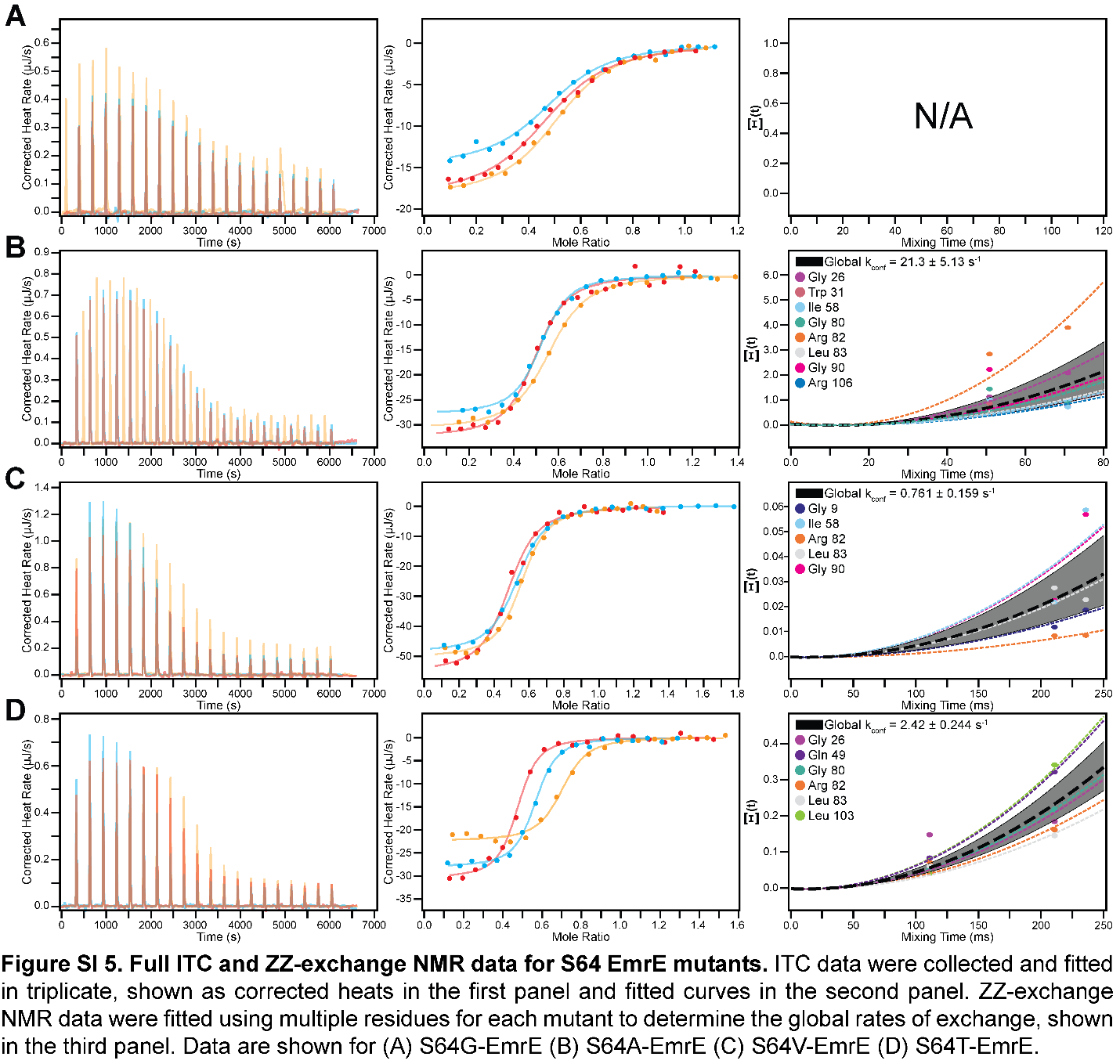

**Figure S4. Full ITC and ZZ-exchange NMR data for S64 EmrE mutants.** ITC data were collected and fitted in triplicate, shown as corrected heats in the first panel and fitted curves in the second panel. ZZ-exchange NMR data were fitted using multiple residues for each mutant to determine the global rates of exchange, shown in the third panel. Data are shown for (A) S64G-EmrE (B) S64A-EmrE (C) S64V-EmrE (D) S64T-EmrE.

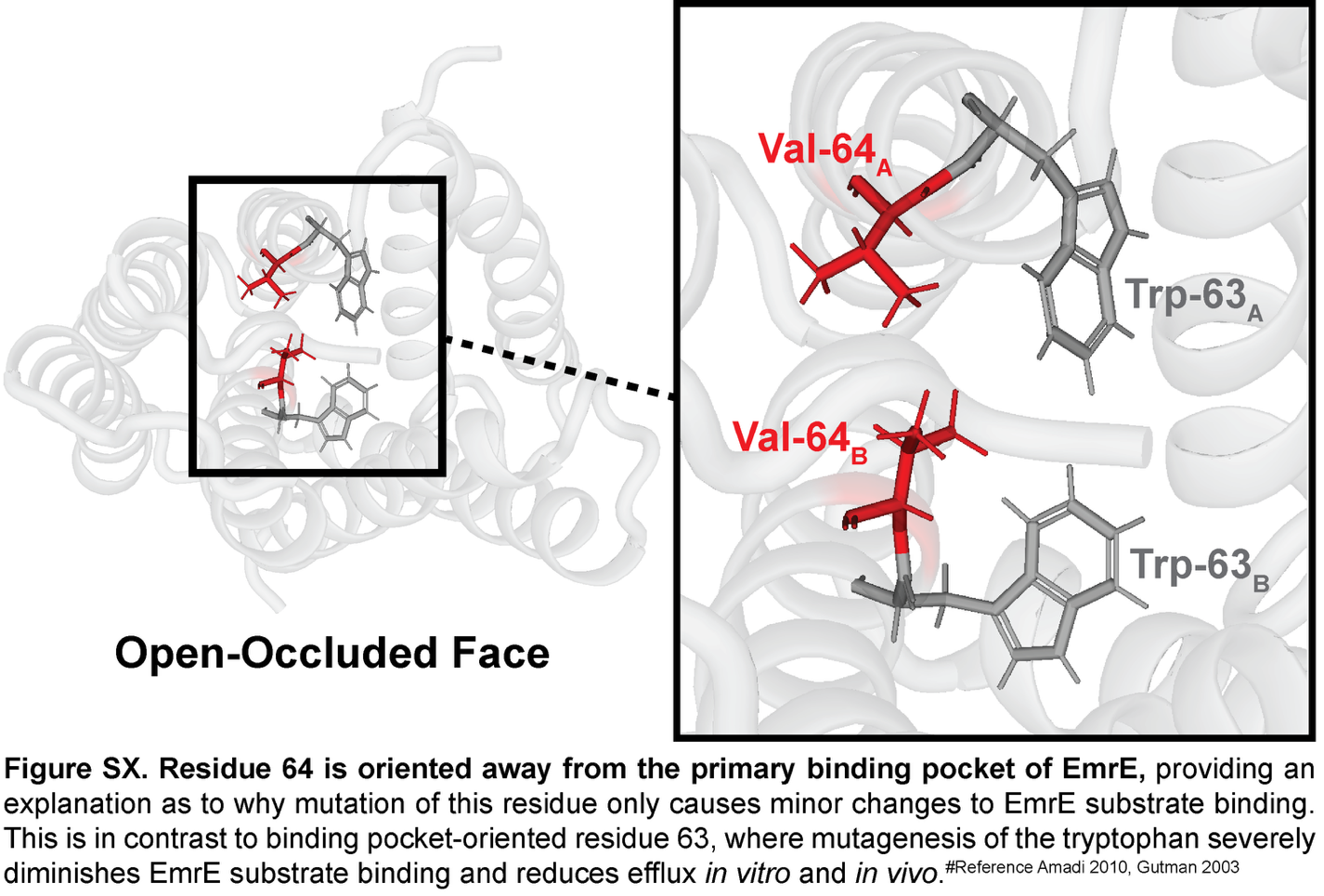

**Figure S5. Residue 64 is oriented away from the primary binding pocket of EmrE**, rationalizing why mutation of this residue only causes minor changes to EmrE substrate binding. This is in contrast to binding pocket-oriented residue 63, where mutagenesis of the tryptophan severely diminishes EmrE substrate binding and reduces efflux *in vitro* and *in vivo*.^1,2^

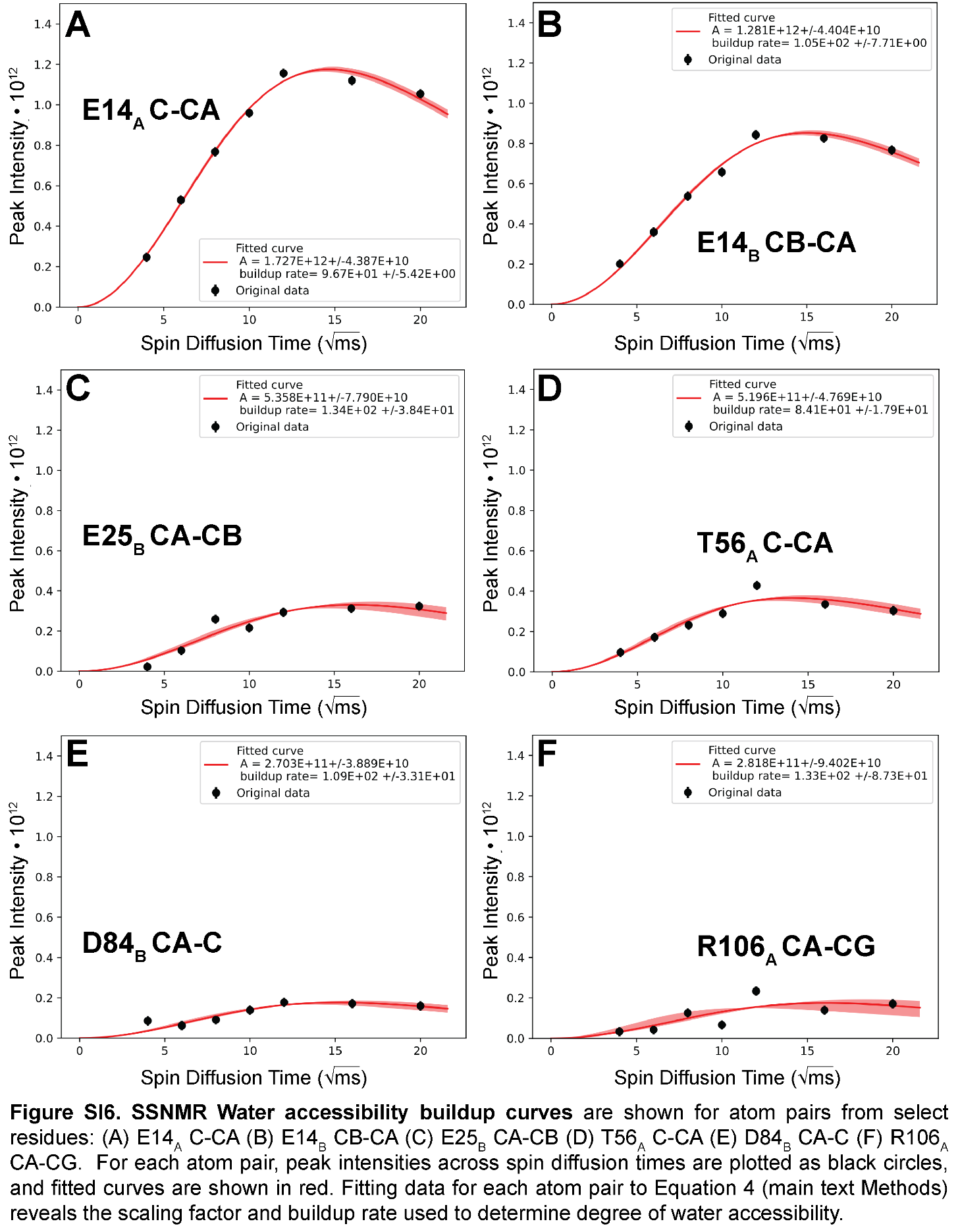

**Figure S6. SSNMR Water accessibility buildup curves** are shown for atom pairs from select residues: (A) E14A C-CA (B) E14B CB-CA (C) E25B CA-CB (D) T56A C-CA (E) D84B CA-C (F) R106A CA-CG. For each atom pair, peak intensities across spin diffusion times are plotted as black circles, and fitted curves are shown in red. Fitting data for each atom pair to Equation 4 (main text Methods) reveals the scaling factor and buildup rate used to determine degree of water accessibility.

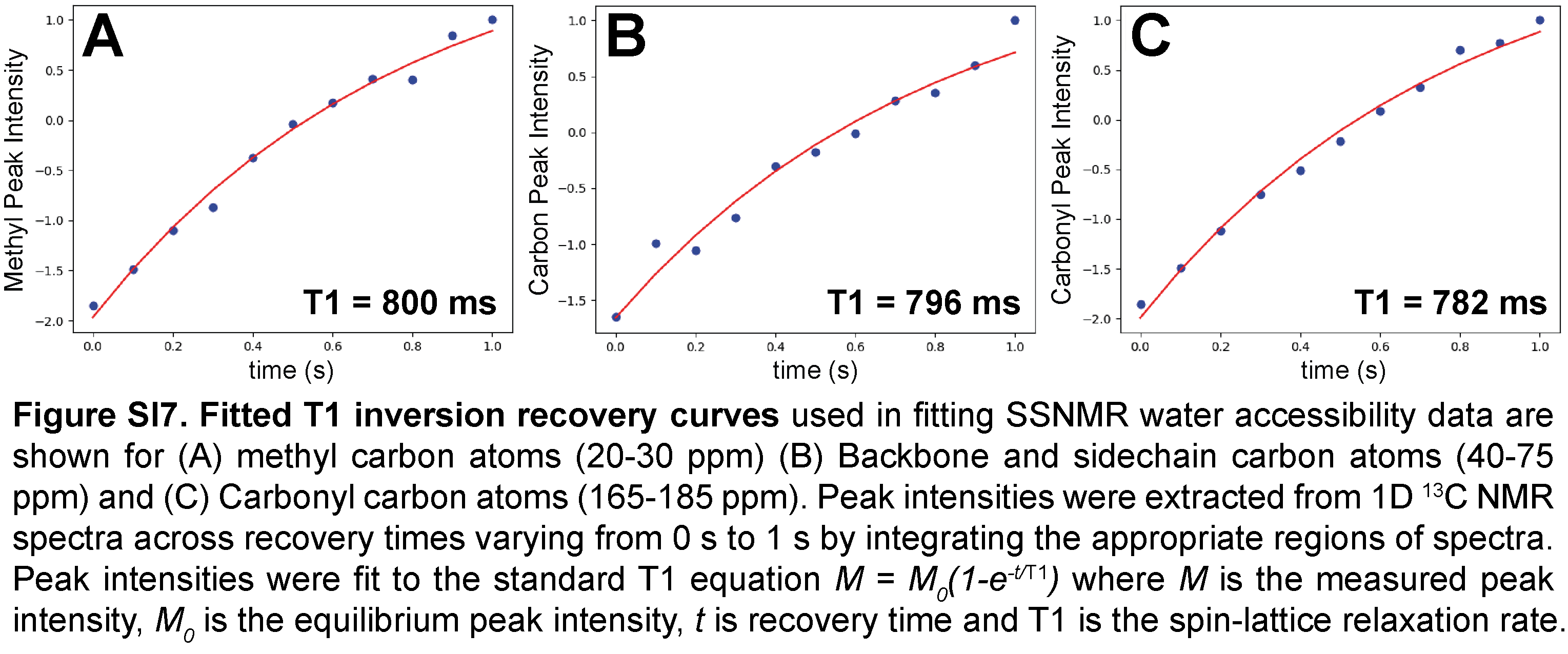

**Figure S7. Fitted T1 inversion recovery curves** used in fitting SSNMR water accessibility data are shown for (A) methyl carbon atoms (20-30 ppm) (B) Backbone and sidechain carbon atoms (40-75 ppm) and (C) Carbonyl carbon atoms (165-185 ppm). Peak intensities were extracted from 1D 13C NMR spectra across recovery times varying from 0 s to 1 s by integrating the appropriate regions of spectra. Peak intensities were fit to the standard T1 equation M = M_0_(1-e^-t/T1^) where M is the measured peak intensity, M_0_ is the equilibrium peak intensity, t is recovery time and T1 is the spin-lattice relaxation rate.

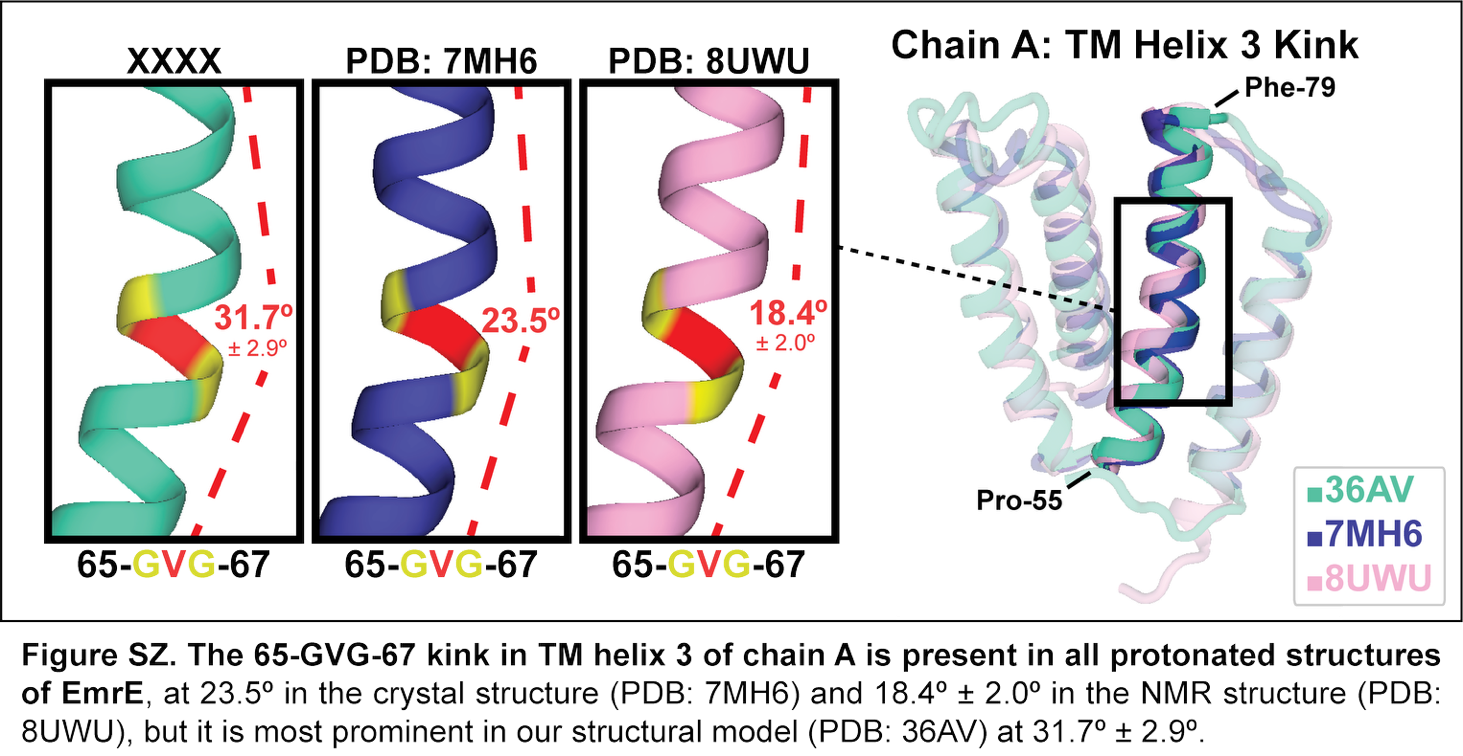

**Figure S8. The 65-GVG-67 kink in TM helix 3 of chain A is present in all protonated structures of EmrE**, at 23.5º in the crystal structure (PDB 7MH6) and 18.4º ± 2.0º in the NMR structure (PDB 8UWU), but it is most prominent in our structural model (PDB 36AV) at 31.7º ± 2.9º.

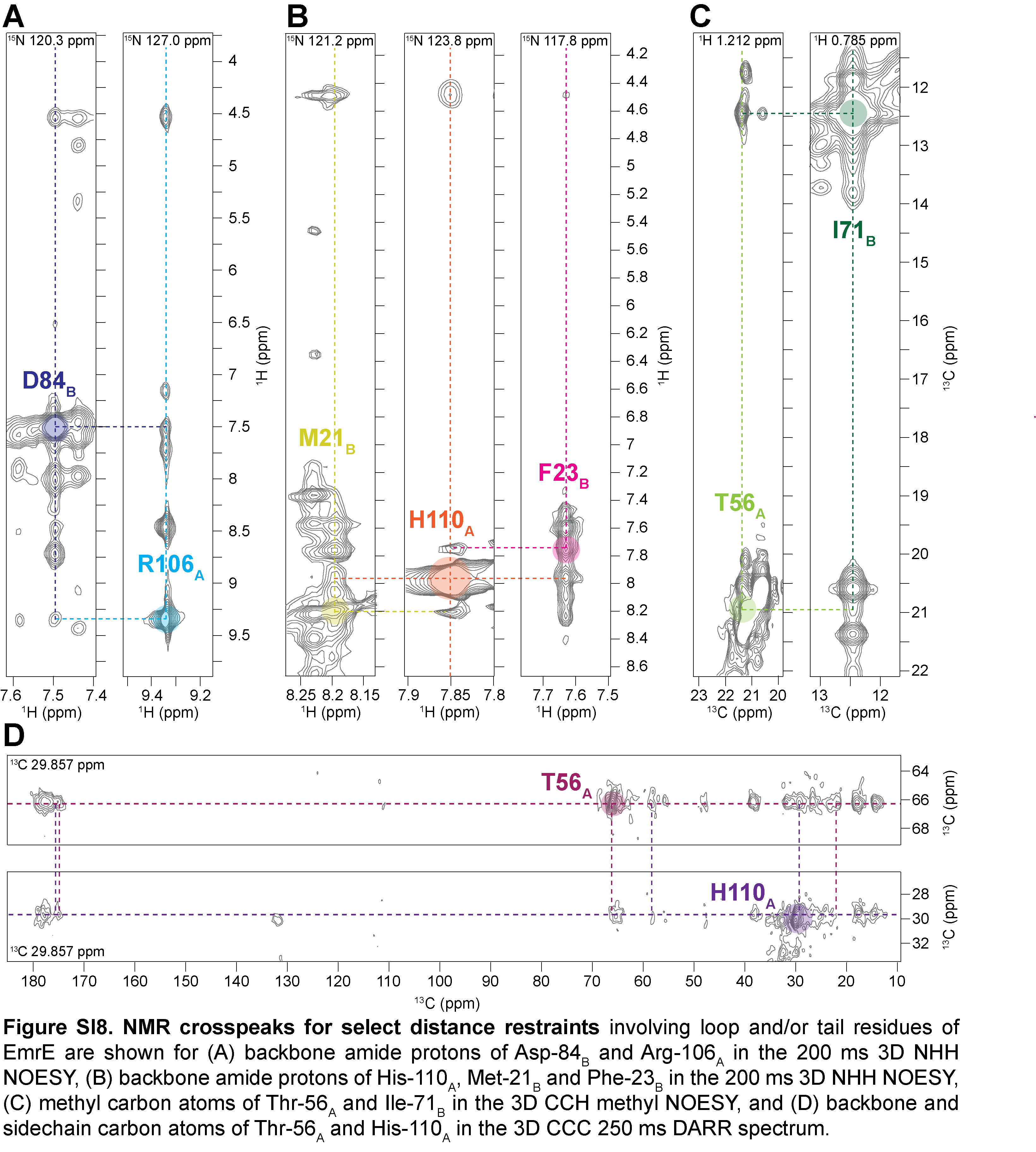

**Figure S9. NMR crosspeaks for select distance restraints involving key loop and/or tail residues** of EmrE are shown for (A) backbone amide protons of Asp-84_B_ and Arg-106_A_ in the 200 ms 3D NHH NOESY, (B) backbone amide protons of His-110_A_, Met-21_B_ and Phe-23_B_ in the 200 ms 3D NHH NOESY, (C) methyl carbon atoms of Thr-56_A_ and Ile-71_B_ in the 3D CCH methyl NOESY, and (D) backbone and sidechain carbon atoms of Thr-56_A_ and His-110_A_ in the 3D CCC 250 ms DARR spectrum.

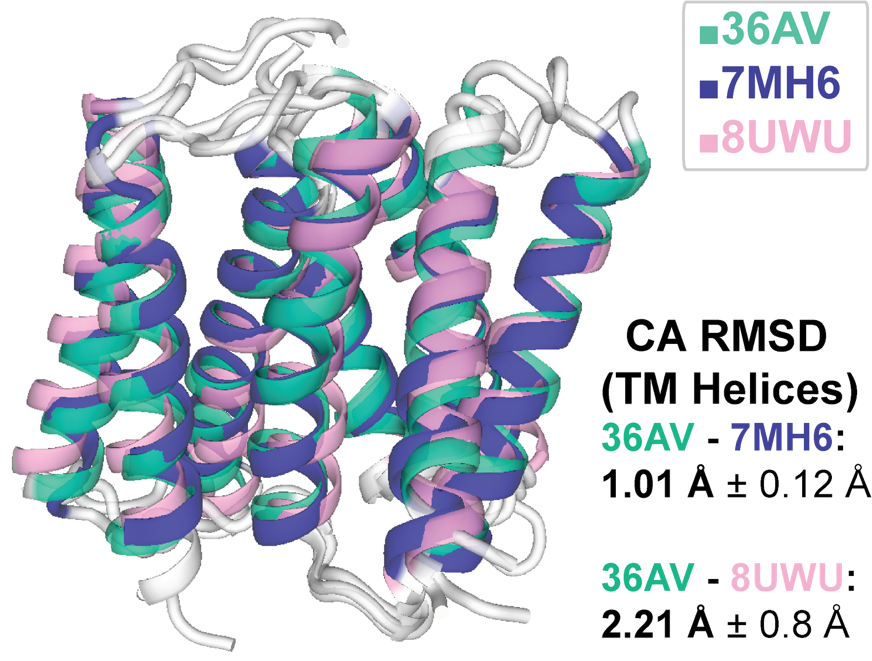

**Figure S10. Backbone CA RMSD for helical regions between our structure and previously published structural models is low**, showing that our NMR restraints within helical regions are largely consistent with previously published models.

| **Spectrum** | **Isotopic labeling pattern** | **Temperature (above or below lipid phase transition)** | **Strong (Å)** | **Medium (Å)** | **Weak (Å)** | **Very Weak (Å)** |
| --- | --- | --- | --- | --- | --- | --- |
| ^13^C-^13^C-^13^C (DARR, t_mix,1_ = 50 ms, t_mix,2_ = 50 ms) | U-^13^C^15^N | below | 4 | 6 | 8 | 10 |
| ^13^C-^13^C-^13^C (DARR, t_mix,1_ = 50 ms, t_mix,2_ = 250 ms) | U-^13^C^15^N | below | 4 | 6 | 8 | 10 |
| ^13^C-^13^C-^13^C (DARR, t_mix,1_ = 150 ms, t_mix,2_ = 600 ms) | sparse | above | 6 | 8 | 10 | 12 |
| NCOCX (DARR, 600 ms) | sparse | below | 4 | 6 | 8 | 10 |

**Table SI 1**. Parameters for each input spectra including isotopic labeling pattern, temperature relative to the lipid phase transition, and binned distances used in the PASD protocol for automated assignment of SSNMR spectra.

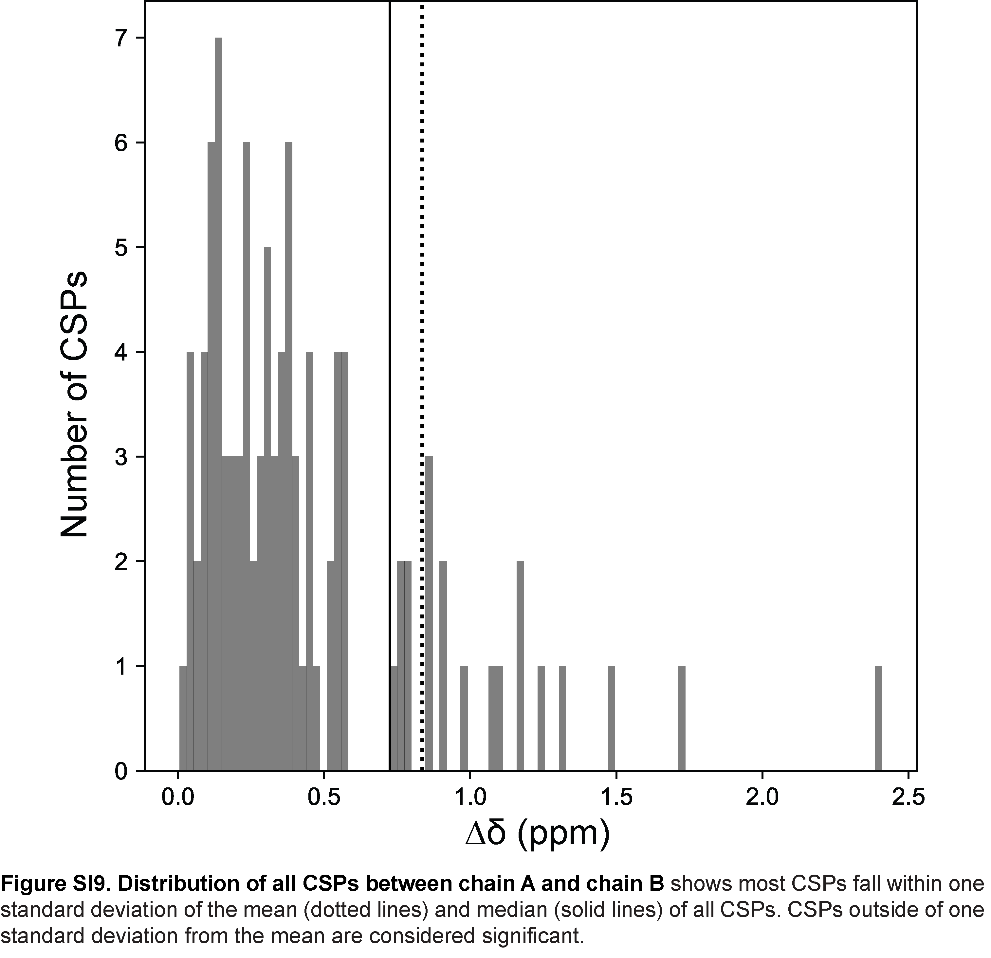

**Figure S11**. **Distribution of all CSPs between chain A and chain B** shows most CSPs fall within one standard deviation of the mean (dotted lines) and median (solid lines) of all CSPs. CSPs outside of one standard deviation from the mean are considered significant.
